# Targeting a shared neoepitope derived from non-canonical translation of c-*MYC* oncogene in cancer cells

**DOI:** 10.1101/2024.05.23.595486

**Authors:** E. Baulu, A. Bolon, E. Etchegaray, F. Raimundo, A. Merienne, J. Martin, J. Grandsire, T. Richard, L. Tonon, C. Dubois, Y. Estornes, R. Boulos, O. Tabone, P. Bonaventura, A. Page, C. Gardet, V. Alcazer, S. Hughes, B. Gillet, N. Gervois, N. Labarrière, Q. Wang, J. Valladeau-Guilemond, N. Chuvin, V. Marcel, J- J. Diaz, S. Depil

## Abstract

Cancer cells rely on alternative modes of translation for protein synthesis, promoting internal ribosome entry site (IRES)-dependent translation of mRNA encoding pro-oncogenic factors. Furthermore, ribosomes translate mRNA with lower fidelity in tumor cells. We proposed that these translational modifications in cancer produce shared tumor-specific epitopes derived from IRES-containing oncogenes. To identify such neoepitopes, we developed an *in silico*-based method that we applied to *c-MYC*. We showed that the non-canonical translation of *c-MYC* mRNA in cancer cells, involving a (+1) ribosomal frameshift, generates a shared neoepitope which induces high-avidity T cells able to kill tumor cells *in vitro* and *in vivo* while sparing normal cells. Our data provide preclinical rationale for developing immunotherapies targeting this *c-MYC*-derived neoepitope and validate a new type of shared translation-associated neoantigens.

## Main Text

The immune response against cancer involves CD8^+^ T cells that recognize tumor epitopes presented on human leucocyte antigen (HLA) class I molecules on the surface of cancer cells. Among the different tumor epitopes, those derived from tumor cell-specific genomic mutations give rise to high-avidity T cells in the absence of thymic selection, which is associated with efficient antitumor cytotoxic properties (*1, 2*). Furthermore, mutation-derived neoepitopes are associated with clinical response to immune checkpoint inhibitors, as evidenced by the high predictive value of the tumor mutational burden (TMB) (*3–8*). Because most of these neoepitopes derive from passenger mutations and are thus patient-specific, targeting this family of tumor-specific epitopes requires personalized and complex approaches (*2*). In addition, many tumors are characterized by a low or moderate TMB, limiting the number of neoepitope candidates (*5*). It is therefore essential to identify new families of tumor-specific antigens shared across patients and tumors for the development of “off-the-shelf” T-cell based immunotherapies (*9*).

Cancer cells are exposed to many stresses (*e*.*g*., hypoxia, nutrient limitation, proteotoxic or genotoxic stresses) and must adapt rapidly to grow and survive. Translation, which represents the last step of gene expression, not only fulfills this requirement but also emerges as a novel source of cancer cell plasticity (*10*). While stress inhibits canonical cap-dependent translation, it promotes alternative modes of translation for protein synthesis, particularly during tumorigenesis. Among these alternative modes of translation, those involving internal ribosome entry sites (IRES) play a key role in cancer since they allow synthesis of proteins with essential pro-oncogenic, anti-apoptotic and survival activities (*10–12*). In tumor cells, ribosomes and transfer RNAs also translate messenger RNAs (mRNAs) with lower fidelity, which induces translational defects including non-canonical initiation, stop codon readthrough or frameshift (*10, 11, 13, 14*). The current concept is that these translational defects, by extending the effective coding range of a given transcriptome, contribute to proteome diversification in cancer cells while constituting a potential “Achilles heel” for the development of new therapies (*15*).

Considering these observations, we hypothesized that the translational defects occurring in cancer cells may produce shared tumor-specific epitopes derived from non-mutated IRES-containing oncogenes. For the proof-of-principle, we selected the *c-MYC* oncogene because (i) it is one of the most frequently amplified and overexpressed oncogenes in tumors, (ii) it is rarely mutated and (iii) its mRNA contains an IRES (*16*).

### *In silico* identification of potential *c-MYC*-derived translational neoepitopes

We first developed an *in silico* method to predict potential neoepitopes derived from the non-canonical translation of *c-MYC* mRNA (Fig. 1A), focusing on HLA-A2, the most common HLA class I allele (*17*). For each of the main *c-MYC* transcript isoforms, we determined all peptide sequences with a size superior or equal to 9 amino acids (AA) that could result from a (+1) or (-1) ribosomal frameshift, stop codon readthrough or any non-canonical initiation or termination of translation. Among those sequences, twenty-two 9-mer peptides were considered as potential strong binder epitopes for HLA-A*02:01 on netMHCpan v4.1. After removing 5 peptides matching a c-MYC protein isoform after alignment with the human proteome, we finally selected 17 potential HLA-A*02:01 epitopes (PR1 to PR17). Two of these predicted peptides corresponded to a potential open reading frame (ORF) within the 5’UTR, 7 to a ribosomal frameshift occurring in the coding sequence (CDS) and 8 to a 3’UTR ORF (fig. S1A). Analysis of mass spectrometry (MS) datasets from The Cancer Genome Atlas (TCGA), the Clinical Proteomic Tumor Analysis Consortium (CPTAC) and Genotype-Tissue Expression (GTEx) showed evidence of translation for PR3 and PR5 peptides in breast and colon cancer samples, but not in normal tissues (fig. S1, A and B). Following confirmation of their effective binding to HLA-A2 molecules in *in vitro* binding assays (fig. S1C), PR3 and PR5 were selected for further analyses.

**Fig. 1.**
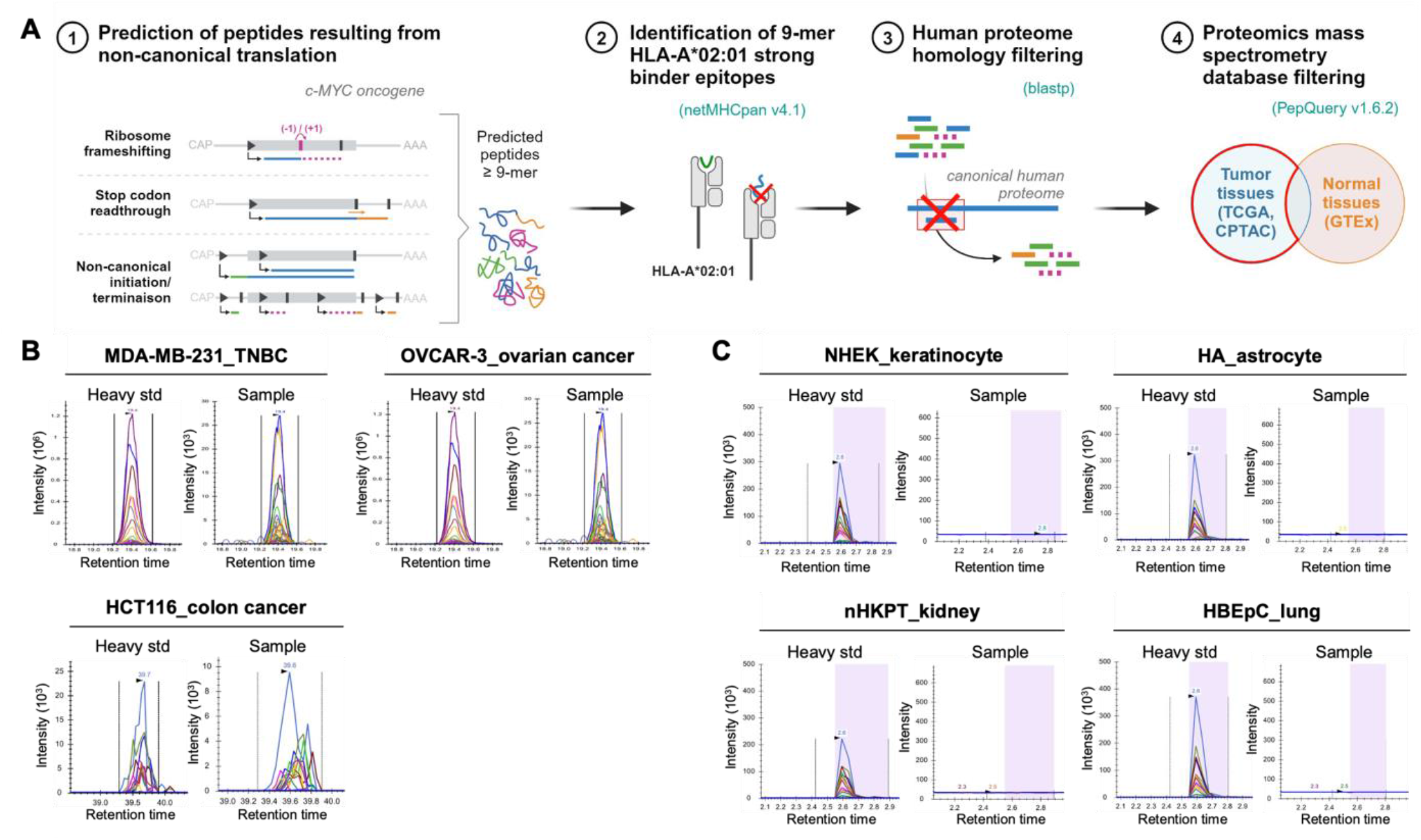
Prediction and validation of the PR3 epitope presentation on tumor cells. (**A**) *In silico* method to predict potential HLA-A*02:01^+^ neoepitopes derived from non-canonical translation of *c-MYC* mRNA. TCGA, The Cancer Genome Atlas; CPTAC, Clinical Proteomic Tumor Analysis Consortium; GTEx, Genotype-Tissue Expression. (**B** and **C**) Valid-NEO transitions of the PR3 peptide (LLLEATANL) on chromatograms, using tumor cell lines (B) or HLA-A2^+^ normal primary cells (C) as samples, and synthetized heavy standard peptides as positive control. std, standard; TNBC, triple-negative breast cancer; NHEK, normal human epithelial keratinocytes; HA, human astrocytes; nHKPT, normal human kidney proximal tubule cells; HBEpC, human bronchial epithelial cells.

### The PR3 peptide is presented exclusively on tumor cells

We assessed the presence of PR3 and PR5 peptides on HLA molecules on the surface of tumor cells. A targeted MS-based method using a highly sensitive technology (*18*) was developed to analyze peptides eluted from HLA molecules. This approach confirmed the presence of PR3 (but not PR5, data not shown) on HLA molecules on the surface of 8 out of 10 HLA-A2-positive tumor cell lines from the NCI-60 panel, representing various types of cancers (triple negative breast cancer, colon, ovarian, non-small cell lung cancer and melanoma) (Fig. 1B and fig. S1D). Importantly, no PR3 peptide was detected on the 4 tested HLA-A2-positive primary normal cells from critical tissues, even with an increased sensitivity provided by a transient boost of the detector (Fig. 1C).

### The PR3 peptide is derived from a (+1) ribosomal frameshift of *c-MYC* mRNA in tumor cells

PR3 was predicted to result from a ribosomal frameshift in the CDS of 9 of the 13 *c-MYC* protein-coding transcript isoforms reported in Ensembl (fig. S2A). However, an additional putative alternatively spliced transcript of *c-MYC* that may contain the PR3 AA sequence in-frame was reported in GenBank (GenBank KC782559.1). Using quantitative RT-PCR, we demonstrated that this alternative transcript is absent from cancer cell lines (fig. S2B). To corroborate this result and further evaluate all *c-MYC* mRNA isoforms not necessarily listed in current databases, we performed long-read RNA sequencing by nanopore in 3 tumor cell lines expressing PR3 on their surface (Fig. 2A). All *c-MYC* transcript isoforms identified using StringTie (*19*) corresponded to transcripts reported in Ensembl and containing PR3 out-of-frame (Fig. 2B). To confirm that the PR3 peptide does not originate from a transcript other than *c-MYC*, potential peptide sequences were predicted from six-frame translation of nanopore long-reads and Illumina short-read assembled-transcripts (Fig. 2A and fig. S2C). The alignment of all the putative peptide sequences containing the PR3 AA sequence confirmed that they all map only to *c-MYC* locus (Fig. 2C and fig. S2D). Likewise, the PR3 AA sequence was searched in NCBI and Ensembl human transcriptome databases by tblastn, and all matches corresponded to *c-MYC* sequences. Finally, PR3 motif did not match any protein in the different human proteome databases. In conclusion, PR3 was only found in *c-MYC* transcripts containing its AA sequence out-of-frame, supporting its translational origin.

**Fig. 2.**
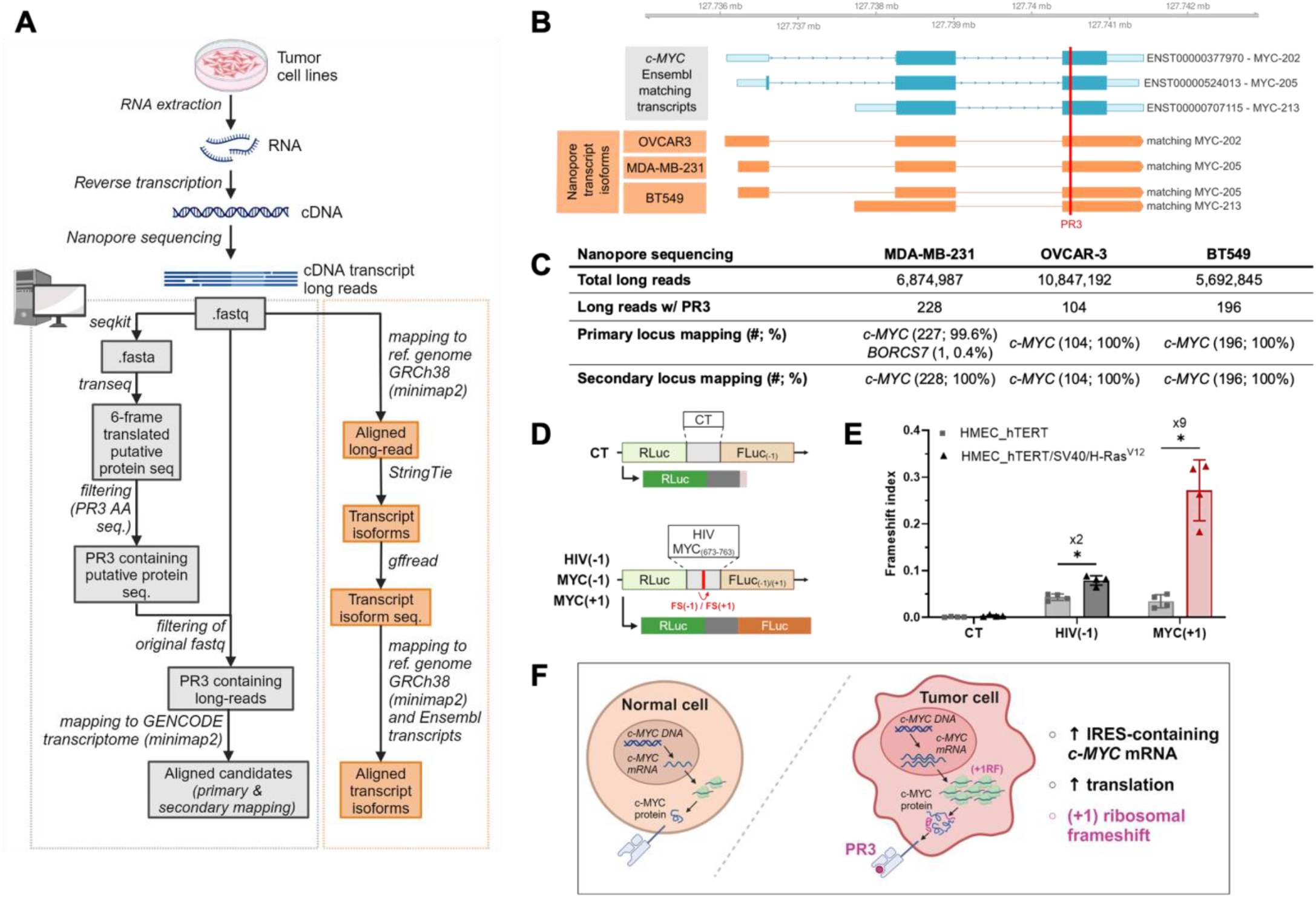
(+1) ribosomal frameshift of *c-MYC* mRNA in tumor cells can generate PR3 epitope. (**A**) Schematic overview of the analysis of the PR3 peptide genomic locus mapping (in grey, left) and the transcript isoform mapping (in orange, right), performed using nanopore sequencing. RNA, ribonucleic acid; cDNA, complementary desoxyribonucleic acid; seq., sequence; ref., reference; AA, amino acid. (**B**) Overview of the nanopore transcript isoforms of MDA-MB-231, OVCAR-3 and BT549 cell lines (in orange) mapped to *c-MYC* locus, compared to matching Ensembl *c-MYC* transcripts (in blue). Transcript’s CDS are in dark blue, untranslated regions in light blue, and the position of the PR3 sequence is indicated with a red line. (**C**) Results of nanopore sequencing and alignment. w/, with; #, number. (**D**) Schematic representation of the bicistronic reporter constructs used for the detection of frameshift events. RLuc is fused to a linker encoding a potential slippery (HIV, MYC_673-763_) or negative control (CT) sequence, upstream of FLuc in -1 or +1 reading frame of RLuc, which is only translated when ribosomal frameshift occurs on the linker sequence. RLuc, renilla luciferase; FLuc, firefly luciferase; CT, control; HIV, human immunodeficiency viruses; FS, frameshift. (**E**) Ratios of the Firefly to Renilla activities (frameshift index) in HMEC_hTERT and HMEC_hTERT/SV40/H-RAS^V12^ cells expressing CT, HIV(-1) or MYC(+1) construct. Results are mean ± s.d. of n=4 independent experiments represented by each dot. * P<0.05, Wilcoxon test. (**F**) Model proposed for the generation of the PR3 epitope in tumor cells. DNA, desoxyribonucleic acid; mRNA, messenger ribonucleic acid; IRES, internal ribosome entry site; RF, ribosomal frameshift.

Next, we developed a bicistronic reporter luciferase assay (*20*) to evaluate both the ability of the *c-MYC* mRNA sequence to induce ribosome slippage and out-of-frame mRNA translation in tumor cells. A *c-MYC* mRNA sequence of 91 nucleotides in length located before the PR3 peptide was inserted between the nucleotide sequences of Renilla (in-frame for a canonical translation) and Firefly (out-of-frame for a translation dependent on a ribosomal frameshift) luciferases (Fig. 2D). Using the HIV (-1) programmed ribosomal frameshift sequence as a positive control, quantification of the activity of Renilla and Firefly luciferase proteins in triple negative breast cancer MDA-MB-231 cells indicated a (+1) ribosomal frameshift in the *c-MYC* mRNA region, while no (-1) frameshift was detected (fig. S2E). Of note, the Renilla/Firefly mRNA ratios were similar in all conditions, confirming that the observed differences were not due to a transcriptional bias (fig. S2F).

To ascertain that this (+1) ribosomal frameshift occurs specifically in tumor cells, we used the same reporter assays in immortalized parental human mammary epithelial cells (HMEC) (transduced with *hTERT*) and transformed HMECs (transduced with *hTERT*, SV40 large-T antigen and *H-rasV12*) (*21*). Whereas the *c-MYC* (+1) ribosomal frameshift index was low in non-tumoral HMECs, a 9-fold increase was observed in transformed cells, much greater than the increase observed for the HIV (-1) ribosomal frameshift, suggesting that the oncogenic process may be at the origin of this (+1) ribosomal frameshift on *c-MYC* mRNA (Fig. 2E and fig. S2G).

Taken together, these results strongly suggest that the tumor-specific PR3 peptide is derived from a (+1) ribosomal frameshift occurring during *c-MYC* mRNA translation in tumor cells (Fig. 2F).

### PR3 is immunogenic and induces high-avidity CD8^+^ T cells

We then sought to determine whether PR3 is immunogenic in cancer patients by studying the presence of specific T cells among tumor-infiltrating lymphocytes (TILs). We analyzed TILs from 22 HLA-A2-positive patients with colon cancer. Due to the low number of T cells available, CD8^+^ T cells were first selected and polyclonally expanded before staining with phycoerythrin (PE) and allophycocyanin (APC)-labelled PR3-HLA-A*02:01 tetramers. Double tetramer staining showed the presence of PR3-specific CD8^+^ T cells in 6 of the 22 (27%) TIL samples tested, suggesting that PR3 is naturally presented and immunogenic in cancer patients (Fig. 3, A and B, and fig. S3A).

**Fig. 3.**
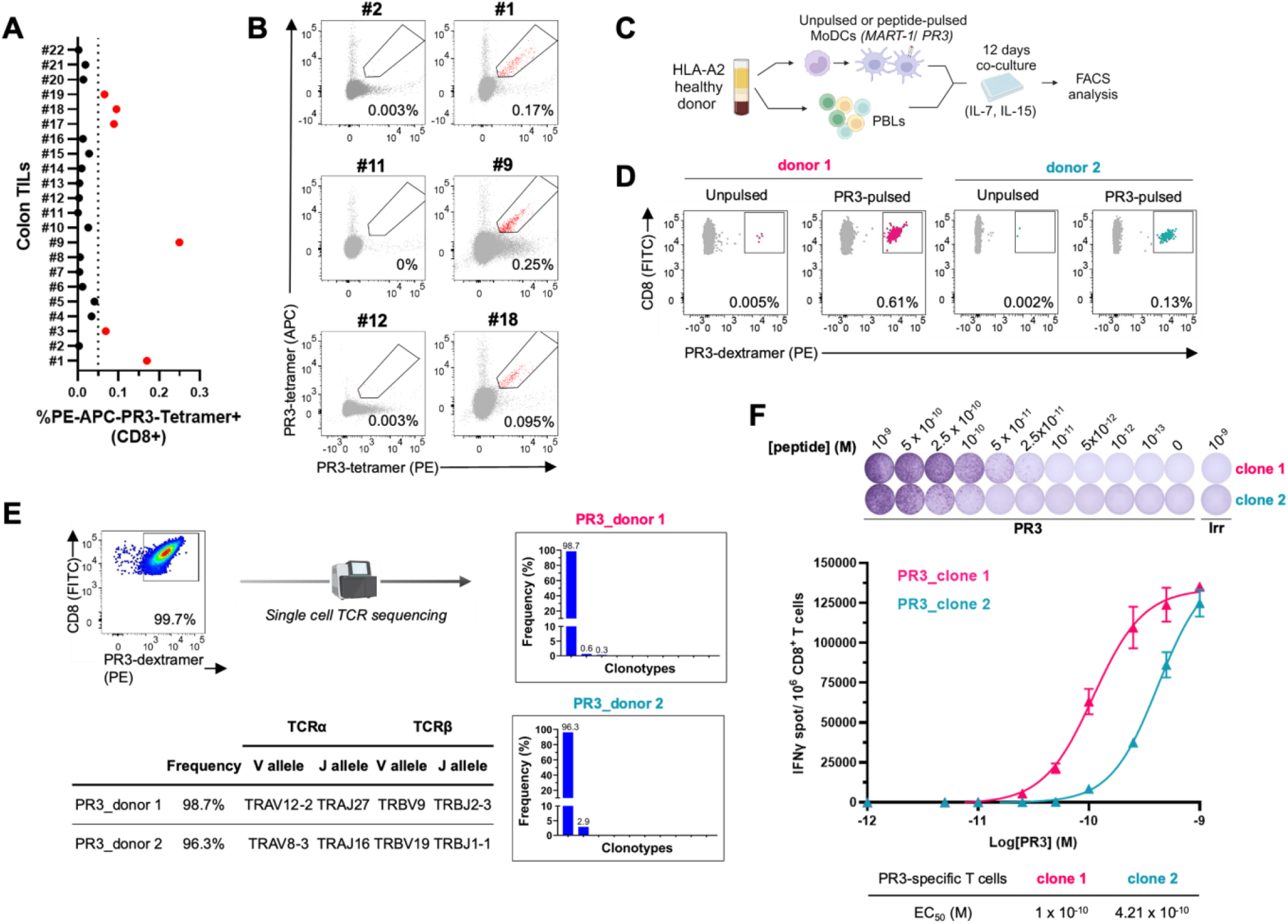
PR3 is immunogenic and induces high-avidity CD8^+^ T cells. (**A**) Quantification of the double PE-/ APC-PR3-tetramer staining of TILs from colon cancer samples gated on CD8^+^ T cells, red dots represent positive quantification. #, number; tetra, tetramer. (**B**) Representative flow cytometry plots and quantification of double PE-/ APC-PR3-tetramer negative (left) and positive (right) staining of TILs from colon cancer samples gated on CD8^+^ T cells. (**C**) Priming protocol. MoDCs, monocyte-derived dendritic cells; PBLs, peripheral blood lymphocytes. (**D**) Representative flow cytometry plots and quantification of PR3-dextramer staining of CD8^+^ T cells from 2 HLA-A2^+^ healthy donors (donor 1 in pink and donor 2 in blue) after 12 days of co-culture with unpulsed or PR3-pulsed MoDCs. (**E**) Clonotype frequencies of PR3-specific CD8^+^ T cells expanded from PR3_donor 1 and PR3_donor 2 using single cell TCR sequencing; and table showing TCRα and TCRβ V and J alleles of the amplified clone. (**F**) Representative IFNγ ELISpot images (top), quantification and nonlinear fit curves of IFNγ spot count per 1 × 10^6^ T cells (bottom) for PR3-specific CD8^+^ T cell clone 1 (pink) and clone 2 (blue), co-cultured with irrelevant-pulsed T2 or T2 pulsed with a limiting dilution of the PR3 peptide. Results are mean ± s.d. of technical duplicates. Data shown are representative of n=4 (PR3_clone 1) independent experiments. EC_50_ is determined using nonlinear fit of mean IFNγ spot count (GraphPad). M, molar mass; [ ], concentration; irr, irrelevant; EC_50_, Half maximal effective concentration.

PR3 immunogenicity was also evaluated by co-culturing peptide-loaded monocyte-derived dendritic cells with T cells from healthy HLA-A2 positive donors (Fig. 3C). After amplification, PR3-specific CD8^+^ T cells were detected by dextramer staining in 4 of the 12 tested donors (Fig. 3D and fig. S3B). Dextramer-stained CD8^+^ T cells obtained from 2 donors were then sorted by flow cytometry and expanded on feeder cells. TCR-sequencing demonstrated the presence of a unique T cell clone after expansion (Fig. 3E). The specificity of both clones was confirmed after co-culture with T2 cells pulsed with PR3 or an irrelevant peptide. Induction of CD137 (4-1BB) (fig. S3C), cytotoxic potential (fig. S3D) and cytokine secretion (IFNγ and TNFα) (fig. S3E) were observed upon stimulation with PR3. To further assess the functional avidity of these T cell clones, we performed an IFNγ ELISpot assay using serial peptide dilutions. Low EC_50_ were observed for the 2 cell clones (1 x 10^-10^ M and 4.2 x 10^-10^ M for clones 1 and 2, respectively) (Fig. 3F). Clone 1 exhibited a particularly high functional avidity with IFNγ secretion detected at a peptide concentration as low as 2.5 x 10^-11^ M, which was associated with efficient killing of tumor cell lines (fig. S3, F and G).

### PR3-specific T cells selectively kill cancer cells

To further assess the antitumor activity of PR3-specific T cells, we transduced primary T cells from HLA-A2-positive donors with the TCR sequenced from clone 1. We initially substituted the human constant regions of TCRα and TCRβ chains by the corresponding murine regions mutated to include an additional disulfide bond to avoid mispairing, as previously described (*22*), with systematic knock-out of the endogenous TCRα/β using CRISPR-Cas9 (fig. S4A). On average, more than 65% of the T cells expressed the transgenic TCR after lentiviral transduction, as shown by double staining with a murine-specific TCRβ antibody and PR3 tetramer (Fig. 4A and fig. S4B). The functionality of TCR-engineered T cells (TCR-T cells) was confirmed after co-culture with T2 cells pulsed with PR3 or an irrelevant peptide, showing specific CD137 (4-1BB) induction (fig. S4C) and secretion of IFNγ, TNFα, IL-2 and granzyme B after activation by PR3-pulsed T2 cells (fig. S4D), using non-transduced T cells as negative controls. ELISpot assay confirmed the high-functional avidity of these TCR-T cells, as evidenced by the detection of specific IFNγ-secreting cells at a PR3 concentration of 5 x 10^-12^ M (fig. S4E). In line with a high functionality, efficient killing of PR3-pulsed T2-cells was still observed at an effector to target cell (E:T) ratio of 1:50 (Fig. 4B).

**Fig. 4.**
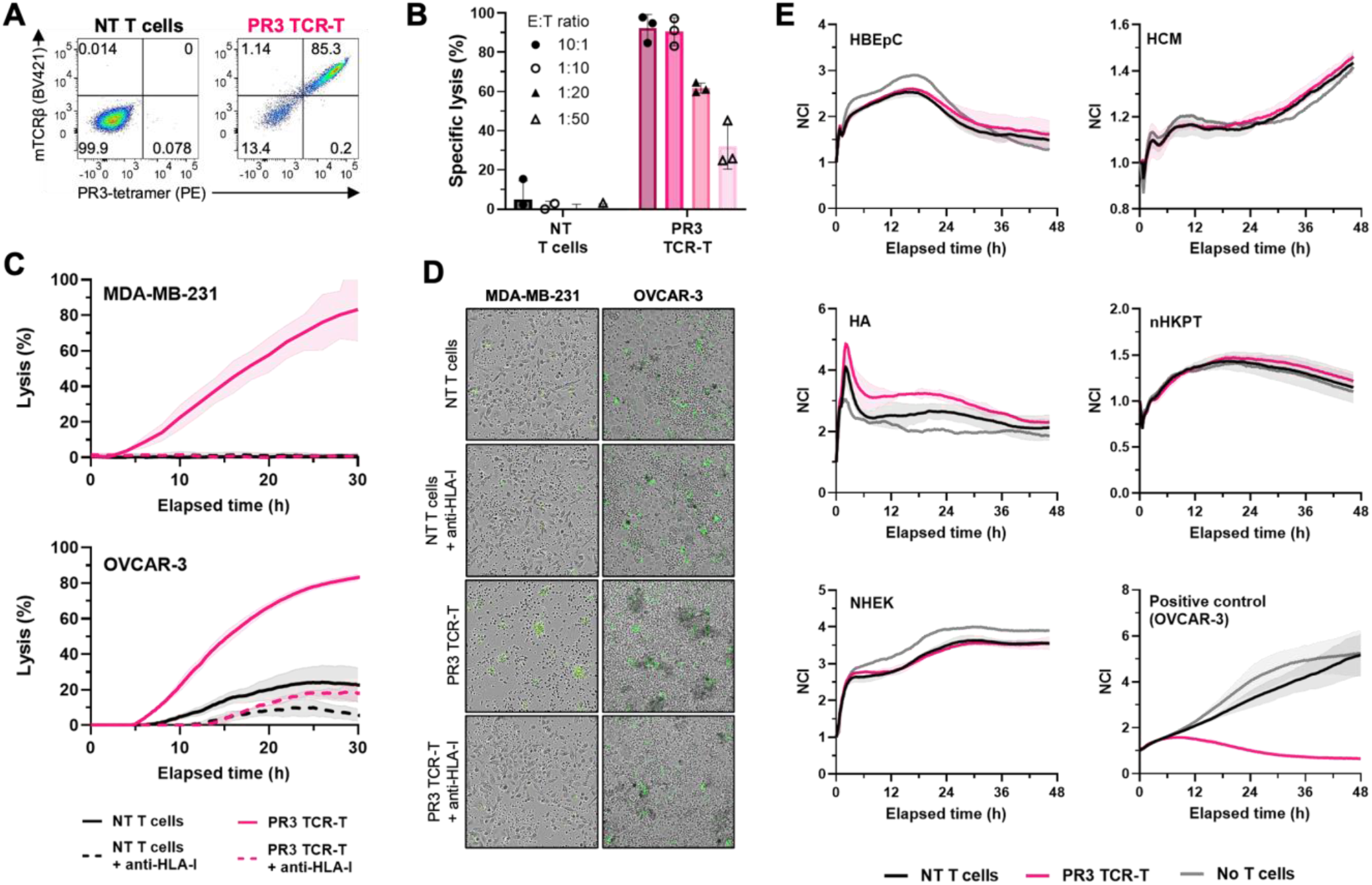
PR3-specific TCR-T cells selectively kill cancer cells. (**A**) Representative flow cytometry plots and quantification of mTCRβ and PR3-tetramer staining of non-transduced (NT) T cells (left) or PR3-specific TCR-T cells (PR3 TCR-T) (right) gated on CD8^+^ T cells. mTCRβ, mouse-TCRβ; NT, non-transduced. (**B**) Specific T cell-induced lysis of PR3-pulsed T2 cells at different effector to target cell ratios. Results are mean ± s.d. percent of n=3 independent experiments represented by each dot. E:T, effector to target cell ratios. NT, non-transduced T cells. (**C**) Tumor cell death induced by PR3-specific TCR-T cells (PR3 TCR-T) at 2:1 effector to target cell ratio. Results are mean ± s.d. of technical triplicates. Data shown are representative of n=2 (MDA-MB-231) independent experiments. NT, non-transduced. (**D**) Representative 10X images of co-cultures presented in (C) at 30h. Dead cells are depicted in green. (**E**) Viability of normal primary cells alone, co-cultured with NT T cells or PR3 TCR-T at 2:1 effector to target cell ratio monitored by normalized cell index through xCELLigence. PR3-expressing tumor cells (OVCAR-3) were used as positive control. Results are mean ± s.d. of technical triplicates. Data shown are representative of n=3 (all normal primary cells) independent experiments. NCI, normalized cell index; HBEpC, human bronchial epithelial cells; HCM, human cardiac myocytes; HA, human astrocytes; nHKPT, normal human kidney proximal tubule cells; NHEK, normal human epithelial keratinocytes; NT, non-transduced.

Next, we assessed the specificity of the selected TCR using AA scanning. TCR-T cells were co-cultured with T2 cells pulsed with glycine (Gly) or alanine (Ala)-substituted peptides to assess the effects of these changes on IFNγ production. The replacement of Ala with Gly at position P5, revealed that this residue was not involved in the interaction of the peptide with the TCR, in agreement with 3D modeling of TCR-peptide-HLA complexes (fig. S5, A, B and E). Ala substitution suggested that Leu in P3 may also be dispensable for this interaction (fig. S5F). To establish the cross-reactivity potential, we thus selected the recognition motif as ‘L-L-x-E-x-T-A-N-L” (“x” corresponding to “any AA”). We then used ScanProsite tool (*23*) to search for this motif in the human proteome. No candidate was found with this approach. To eliminate potential cross-reactive epitopes with different AA at anchorage positions P2 and P9, which may be omitted after classical Gly and Ala scanning, we extended the motif to “L-x-x-E-x-T-A-N-x”. Eight candidates were identified but none was predicted to bind HLA-A*02:01. Taken together, these analyses confirmed the specificity of this TCR for PR3.

TCR-T cells were then co-cultured with cancer cell lines expressing PR3 in immunopeptidomics and tumor cell killing was measured in real time by IncuCyte or xCELLigence. TCR-T cells efficiently killed tumor cells, while no cytotoxicity was observed with non-transduced T cells (Fig. 4C and fig. S4F). Killing was blocked by anti-HLA-I antibody, confirming that the observed cytotoxicity is HLA-dependent (Fig. 4C). Imaging showed tumor cell death and T cell clustering when PR3 TCR-T cells were co-cultured with cancer cells (Fig. 4D). In agreement with immunopeptidomic analyses showing the absence of PR3 on normal cells, no activation and no cytotoxicity were observed when TCR-T cells were co-cultured with HLA-A2-positive primary cells from normal critical tissues (Fig. 4E and fig. S4, G and H).

The *in vivo* anti-tumor efficacy of PR3-specific TCR-T cells was then investigated. We used an avian embryo model to establish 3D tumors within a few days in embryonic regions providing a microenvironment supportive of tumor formation (*24*) (Fig. 5A). PR3-specific TCR-T cells co-transplanted with the triple negative breast cancer cell line MDA-MB-321 or the ovarian cancer cell line OVCAR3 exhibited a strong anti-tumoral activity, inducing a highly significant decrease in tumor volume compared to non-transduced T cells. Indeed, a mean decrease of 28% (P<0.05) and 59% (P<0.001) in normalized tumor volumes was observed in embryos transplanted with TCR-T cells compared to the non-transduced T cells, for MDA-MB-231 and OVCAR-3, respectively (Fig. 5, B to D, and fig. S6, A to D).

**Fig. 5.**
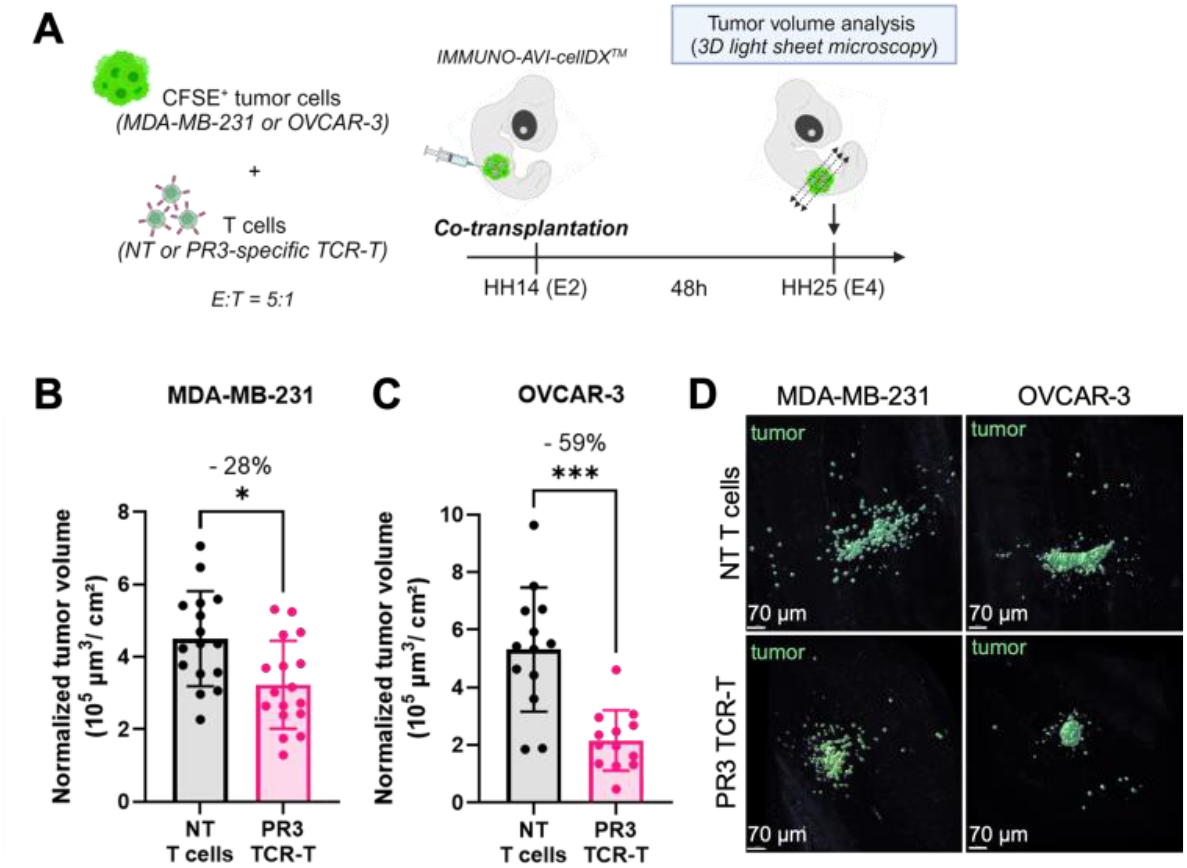
PR3-specific TCR-T cells elicit *in vivo* anti-tumor effects. (**A**) Schematic representation of the *in vivo* experiments. E:T: effector to target cell ratio; NT: non-transduced; HH, Hamburger-Hamilton stage; E, embryonic day. (**B** and **C**) Quantification of MDA-MB-231 (B) or OVCAR-3 (C) tumor volumes using 3D light sheet microscopy of fluorescent target cells, after 48h co-engraftment with CD8^+^ NT T cells or PR3 TCR-T at 5:1 effector to target cell ratio. Tumor volumes are normalized against body surface area. Results are mean ± s.d. of 33 embryos (NT=16; PR3 TCR-T =17) (B) and 26 embryos (NT=13; PR3 TCR-T=13) (C) represented by each dot. *P<0.05; ***P<0.001, Wilcoxon tests. NT, non-transduced. (**D**) Representative images at 48h of 3D views (light-sheet imaging) HH25 chick embryos co-engrafted with CFSE-labelled MDA-MB-231 (left) or OVCAR-3 (right) and NT T cells (top) or PR3 TCR-T (bottom).

Altogether, these data demonstrate that PR3-specific T cells specifically recognize and kill tumor cells, both *in vitro* and *in vivo*, while sparing normal cells.

## Discussion

These results support our main hypothesis that non-canonical translation events associated with the tumorigenic process generate neoepitopes from IRES-containing oncogenes (*10–12*). Using an in-house bioinformatic approach focusing on non-canonical translation, we identified and validated the translation-associated neoepitope PR3, derived from the *c-MYC* mRNA oncogene, and showed that it induced an efficient and specific antitumor T cell response.

The PR3 AA sequence originates from a predicted ribosomal frameshift in the CDS of *c-MYC* mRNAs. We confirmed that the PR3 peptide arose from a translational event, as its sequence was only found in reported *c-MYC* transcript isoforms containing PR3 out-of-frame. Furthermore, the results of the reporter assays led to the conclusion that PR3 most likely derives from a (+1) ribosomal frameshift occurring during *c-MYC* mRNA translation in tumor cells. Programmed ribosomal frameshifting (PRF) is a common mechanism in viruses to produce several essential proteins from a single viral RNA. In this case, PRF is often initiated by an RNA secondary structure and a slippery nucleotide sequence, such as for the (-1) PRF motif of HIV used here as a positive control (*25, 26*). In cancer cells, (+1) ribosomal frameshifting has been described in response to AA shortage; a phenomenon called “sloppiness” (*27*). The ribosomal frameshifting induced by the absence of tryptophan was shown to be associated with the generation and presentation of aberrant peptides able to induce T cell responses (*28*). Of note, bioinformatics analyses did not identify any specific mRNA structure nor motif associated with this ribosomal frameshift (*28*). Interestingly, a link was reported between the levels of sloppiness and oncogenic mutations, particularly in the mitogen-activated protein kinase RAS-RAF signaling pathway (*27*). Similarly, we observed a significant increase in the *c-MYC* (+1) ribosomal frameshift index in *H-rasV12* transformed HMECs compared to immortalized HMECs, supporting the role of the RAS oncogenic pathway in (+1) ribosomal frameshifting.

The translation-associated neoepitope PR3 was specific to cancer cells in immunopeptidomics. We propose that the combination of the 3 following factors: (i) increase in *c-MYC* mRNA levels due to genomic alterations, (ii) increase in translation of IRES-containing mRNAs, including *c-MYC*, and (iii) induction of (+1) ribosomal frameshifting during tumorigenesis, concur to the generation of this *c-MYC*-derived neopeptide only in cancer cells (Fig. 2F). Interestingly, *c-MYC* is one of the most frequently expressed oncogenes and is rarely mutated in solid tumors (*16*). Targeting a shared translational neoepitope derived from *c-MYC* would thus be of a great therapeutic interest. We showed that CD8^+^ T cells engineered with a PR3-specific TCR selectively killed cancer cells. Further studies are required to evaluate T cell constructs optimized in terms of persistence and resistance to exhaustion in different *in vivo* models (*29*). Nevertheless, these data provide the preclinical rationale for developing T-cell based immunotherapies targeting PR3, such as TCR-T cell therapies or bispecific antibodies specific to PR3-HLA-A2 complex (*30*).

We anticipate that our *in silico* method could be applied to identify additional neoepitopes derived from other IRES-containing oncogenes or survival factors, and thus shared across patients and tumors. It would also be interesting to extend the survey to other frequent HLA alleles. This type of “Translation-Associated NeoAntigens (TANAs)” may represent an alternative to mutation-associated neoantigens (MANAs), especially for the treatment of “cold tumors” that have a low mutational burden.

## Acknowledgments

HMEC_hTERT and HMEC_hTERT/SV40/H-Ras^V12^ cell lines, and pCMVdeltaR8.91 and phCMVG-VSVG packaging plasmids were a gift from Dr. AP Morel (Cancer Research Center of Lyon (CRCL), Lyon, France). The illustrations were created with https://BioRender.com.

## Funding

This project, accredited by Lyonbiopôle Auvergne-Rhône-Alpes, has been carried out thanks to the support of the Cancéropôle CLARA and La Région Auvergne-Rhône-Alpes as part of the Proof-of-Concept program (project NEOCELL), and by the Ligue contre le Cancer. EB received a CIFRE from ErVimmune.

## Author contributions

Conceptualization: EB, NC, VM, JJD, SD. Methodology: EB, AB, EE, FR, JM, LT, CD, YE, RB, OT, PB, VA, NG, QW, NC, VM, SD. Investigation: EB, AB, EE, FR, AM, JM, JG, TR, LT, CD, RB, CG, SH, BG, QW. Supervision: EB, YE, RB, OT, PB, AP, NG, NL, JVG, NC, VM, JJD, SD. Writing – original draft: EB, EE, VM, JJD, SD. Writing – review & editing: EB, EE, JVG, NC, VM, JJD, SD

## Competing interests

EB, NC, VM, JJD and SD are co-inventors on a patent application filed on the subject matter of this study (WO/2024/062113).

## Data and materials availability

All data are available in the main text or the supplementary materials.

## Supplementary Materials

Materials and Methods

Figs. S1 to S7

Tables S1 to S3

References (31–63)

## Notes

### Competing Interest Statement

The authors have declared no competing interest.

