## Supplementary data for "Targeting a shared neoepitope derived from non-canonical translation of c-*MYC* oncogene in cancer cells"

**The PDF file includes:**

Materials and Methods
Figs. S1 to S7
Tables S1 to S3

### Materials and Methods

In silico neoepitope prediction

On the 2020/11/20, all *c-MYC* transcripts (n=7) were identified using Ensembl (*31*). For each transcript, all open-reading frames (ORF) were predicted from AUG or non-AUG-start codons to the first or the second stop codon (*i.e.*, to identify potential stop codon readthrough), in all mRNAs annotated sequences (*i.e.*, 5'-end to 3'-end, including UTR regions to encompass potential non-canonical initiation or termination of translation). All predicted ORFs were translated in the 3 frames (*i.e.*, to cover potential translational frameshift). Among those putative peptide sequences, 9-mer strong binder epitopes for HLA-A*02:01 were predicted using netMHCpan v4.1 (*32*). Any potential epitope with a perfect sequence homology to the human proteome was removed from the previous selection (blastp, refseq_protein database). Predicted 9-mer peptides were searched in peptidome proteomic datasets using Pepquery (v.1.6.2.0) (*33*) on the 2020/11/20, as previously described (*34*). Tandem mass spectrometry (MS/MS) datasets from colon cancer (CPTAC colon (*35*) and TCGA colon (*36, 37*)), breast cancer (CPTAC breast (*38*) and TCGA breast (*39*)) and normal tissues (GTEx (*40*)) were analyzed. Briefly, raw MS/MS datasets were downloaded and retrieved MS/MS spectra were converted to MGF format using msconvert from proteowizard (*41*). The data were processed using the following command line in Pepquery: java -Xmx10G -jar pepquery.jar -fixMod 6,62,108 -varMod 117 -maxVar 3 -c 1 -tol 10 -tolu ppm -minScore 12 -e 1 -um -hc TRUE -n 1000 -itol 0.05 -m 1 cpu 12 -pep ${peptides_list} -db ${Reference_database} -ms ${MS_database} -o ${output_directory}. A peptide was selected when the identification was "confident" according to Pepquery validation steps (*33*).

Peptides

HPLC grade 9-mer peptides (> 90%) were manufactured by Thermo Fisher Scientific. Lyophylized powder was resuspended in 1% DMSO (Sigma-Aldrich) distilled water.

Binding affinity analysis

The 2 selected (PR3 and PR5) and positive control peptides were synthetized to perform a high throughput quantitative binding assay to MHC allele HLA-A2, using ProImmune's Class I REVEAL® Rapid Epitope Discovery system as previously described (*42*).

Biological samples and TILs preparation

PBMCs were isolated from healthy donor blood samples ("Etablissement Français du Sang", Lyon, France) using Ficoll density gradient centrifugation (Eurobio). Colon cancer samples (n=22) were provided by the tissue bank of Nantes University Hospital (CHU, Nantes, France; French agreement number DC-2014-2206), with the approval of the ethics commit (CPP Ouest IV - Nantes) and patient written informed consent, in accordance with the Declaration of Helsinki and the guidelines of the French Ethics Committee for Human Tissue Research. TILs were obtained by culturing small tumor fragments (~1 $mm^3$) in RPMI1640 (Gibco) supplemented with 8% AB human serum (Dutscher), 2 mM GlutaMAX (Gibco), 1% penicillin-streptomycin (PS, Gibco), 1 µg/mL amphotericin B (Sigma-Aldrich), 0.1 mg/mL gentamycin (Sigma-Aldrich) and 150 U/mL human recombinant interleukine-2 (hrIL-2, Proleukin, Novartis Pharma) for ~3 weeks. $CD8^+$ TILs were collected by positive magnetic cell sorting using the REAlease CD8 MicroBead kit (Miltenyi) according to the manufacturer's instructions. CD8+ TILs were expanded on irradiated allogeneic feeder cells (PBMCs and Epstein-Barr virus-immortalized B cells) with 1 µg/mL phytohemagglutinin-L (PHA, Sigma-Aldrich) and 150 U/mL hrIL-2, as previously described (*43*).

Cell lines and primary cells

MDA-MB-231 (Cat#ACC 732), HCT116 (Cat#ACC 581) and T2 (Cat#ACC 598) cells were purchased from DSMZ and cultured according to manufacturer's instructions except for MDA-MB-231, which was cultured in DMEM (Gibco) with 10% fetal bovine serum (FBS, Eurobio Scientific). OVCAR-3 (Cat#HTB-161), BT549 (Cat#HTB-122) and 293T (Cat#CRL-11268) cells were purchased from ATCC and cultured according to manufacturer's instructions. HMEC_hTERT and HMEC_hTERT/SV40/H-Ras$^{V12}$ (kind gift of Dr. AP Morel (*21*)) cells were cultured in DMEM/F-12 (Gibco) with 10% FBS, 0.01 µg/ml hEGF (Sigma-Aldrich), 0.5 µg/ml hydrocortisone (Sigma-Aldrich) and 280 mU/ml of human insulin (Humalog, Eli Lilly). For all tumor cell lines, 1% penicillin-streptomycin (PS, Gibco) was added to the culture medium. HLA-A2+ normal primary cells HCM (Cat#C-12810, lot#463Z016.2, female donor DN000249), HBEpC (Cat#C-12640, lot#469Z016, female donor DN000331) and NHEK (Cat#C-12003, lot#467Z005.1, female donor DN000068) were purchased from Promocell, nHKPT cells (Cat#3253021, lot#001C, male donor) from Tebubio and HA cells (Cat#1800-SC, lot#35382) from ScienCell. All human normal primary cells were cultured following the supplier's recommendations. All cells were grown at 37°C in 5% CO2 with humidification. BT549, HCT116, SW-620, SK-MEL5, NCI-H522, OVCAR-5, MCF-7 and COLO205 tumor cell lines (NCI-60 panel) for MS-immunopeptidomics analysis were provided by Complete Omics Inc. MDA-MB-231 cells expressing the Histone 2B-mcherry fusion protein (mCherry nuclear marker, mCherry_MDA-MB-231) were generated by retroviral transduction. Briefly, 293T cells were seeded at 0.8 x $10^6$ cells per 100-mm² and were transfected 3 days later with the following plasmid mix: pHIVgag-pol : pMDG : pH2B_mCherry plasmids (kindly provided by Dr. G. Ichim, Centre de Recherche en Cancérologie de Lyon), at a 1:1.5:1.2 ratio, using the Calcium Phosphate transfection kit (Invitrogen) according to the manufacturer's protocol. 48 h and 72 h after transfection, viral supernatants were collected, pooled and centrifuged for 16h at 3,000g and 4°C. The next day, H2B_mCherry retroviruses were added to MDA-MB-231 cells, plated the day before at 150,000 cells per well in 6-well plate, in presence of 5 µg/ml polybrene. After a 48h incubation, media was renewed and mCherry expressing MDA-MB-231 cells were amplified and sorted using BD FACSAria Cell Sorter (BD Bioscience) until reaching > 90% purity.

Epitope validation by MS-immunopeptidomics

Epitope validation was performed by Complete Omics Inc. (Maryland, USA) according to the method previously described (*18*) with additional modifications. Briefly, 20M cells were lysed and peptide-HLA complexes were isolated using an in-house packed Valid-NEO neoantigen enrichment column preloaded with anti-human Pan-HLA antibodies modified with conjugated chemical moieties designed to increase the binding efficiency between the column matrix and HLA neoantigen complex molecules. The MaxRec version 5 technology was used to increase the sensitivity of the detection by preventing sample loss. After elution, dissociation, filtration, and cleanup, antigen peptides were lyophilized before further analysis. Detection parameters were examined and curated through a Valid-NEO method builder bioinformatics pipeline developed by Complete Omics Inc. Precursor or fragmented ions with excessive noise due to environment or coelution with impurities were excluded from the detection. To boost the detectability, a series of recursive optimizations for the significant ions was conducted.

Database search analyses

To verify PR3 origin, PR3 amino-acid (AA) motif was searched in human transcriptome using tblastn against Ensembl Human GRCh38 cDNAs (transcripts/splice variants) and NCBI Transcript Reference Sequences databases. PR3 AA motif was also searched in human proteome using blastp analyses against Ensembl, NCBI refseq_protein, UniProtKB/SwissProt and PDB human protein databases. Blast analyses were run on v2.15.0, with parameters adjusted for short query sequences (maximum E-value 100000, word size of 3, PAM30 scoring matrix) (*44, 45*).

Data generation and analysis of the nanopore RNA sequencing

Total RNA was extracted with the RNeasy Plus Mini Kit (Qiagen) according to the manufacturer's protocol. RNAs were quantified by Qubit 4.0 (RNA assay kit, Thermofisher Scientific) and qualified with Tapestation 4150 (RNA ScreenTape, Agilent). cDNA production and sequencing were performed using the Direct cDNA Sequencing V14 protocol (SQK-LSK114) of Oxford Nanopore Technologies (ONT). 220-500 ng cDNA were produced and used for each library construction. Libraries were sequenced separately with R10.4.1 flow cells (FLO-MIN114, ONT) using Mk1C devices and MinKNOW v23.04.8 or v23.07.12. Pod5 files were reanalyzed post-sequencing with Super Accurate basecalling performed with MinKNOW P2 Solo v23.07.12 and Guppy v7.1.4. allowing read splitting. Read quality control was conducted using pycoQC (*46*). The Pass filter threshold, separating failed reads from passed reads, was set at Q10.

For the splice variant search, sequenced long reads fastq were converted to fasta with seqkit v4-r122 (*47*). Long read sequences were compared to the transcript sequences reported to encode the *c-MYC* splice variant (GenBank: KC782559.1; UniProtKB: T2B5D4_HUMAN), with a tblastn (v2.14.0+) analysis using blast+ package (*48*).

For the genomic locus mapping of the PR3 peptide, fasta reads were translated in the 6 frames using transeq (*49*) (version EMBOSS:6.6.0.0). The putative translated peptide sequences were filtered to retain only those containing the PR3 AA sequence. Long reads corresponding to the putative PR3-containing peptides were then mapped to the reference GENCODE v33 transcriptome using minimap2 (*50*) (using splice option, v2.26-r1175). Both the primary and secondary mapping were assessed to identify the mapped contig.

For transcripts isoforms analysis, nanopore long reads were mapped to the human reference genome GRCh38 with minimap2 (using splice option, v2.26-r1175) (*50*). Then, transcript isoforms were identified with StringTie version 2.2.1 using long read option (*19*). Transcript isoform sequences were retrieved with gffread (v0.12.8) using the human reference genome GRCh38 (*51*). Then the transcript isoforms were mapped to the reference GRCh38 with minimap2 (using splice option, v2.26-r1175) and compared to *c-MYC* transcript isoforms described in Ensembl using clustal omega (v1.2.4) (*52*).

Data generation and analysis of the Illumina RNA sequencing

Total RNA was extracted, quantified, and qualified as for nanopore RNA sequencing data generation. RNAs were purified using the NEBNext® Poly(A) mRNA Magnetic Isolation Module protocol (New England Biolabs, E7490L). Indexed libraries were constructed following the Lexogen CORALL Total RNA-Seq Library Prep Kit V2 with UDI 12 nt Set B1 protocol (Clinisciences, 181.96). Library quality was verified with Qubit 4.0 (RNA assay kit, Thermofisher Scientific) and Tapestation 4150 (RNA ScreenTape, Agilent). The sequencing was performed on an Illumina NextSeq 500, using a NextSeq High Output Flow Cell v2.5. Read quality control was conducted using FastQC and MultiQC and no reads were discarded (*53*).

For the genomic locus mapping of the PR3 peptide, short reads were mapped to the human reference genome GRCh38 with STAR (v2.7.10b), using reference GENCODE v33 transcriptome annotation file to improve accuracy of the mapping, and filtering reads with more than 10 multimapping and reads with more than 7 mismatches (*54*). Transcripts were assembled from aligned files using StringTie version 2.2.1 (*55*). Assembled transcript sequences were then retrieved using gffread (v0.12.8) and were translated in the 6 frames using transeq (version EMBOSS:6.6.0.0). The putative translated transcripts were filtered out to keep only those containing PR3 AA sequence. PR3-containing transcript sequences were then mapped to the reference genome GRCh38 with minimap2 (using splice option, v2.26-r1175).

<u>Bicistronic reporter constructs</u>

To investigate ribosomal frameshift, four bicistronic reporter plasmids (pTRIP_CT, pTRIP_HIV(-1), pTRIP_MYC(-1) and pTRIP_MYC(+1)) were constructed using pTRIP vector as backbone (GenScript). Each construct is composed of (i) a first cistron, encoding the Renilla Luciferase (RLuc) reporter protein and whose translation is dependent of the cap, (ii) a linker sequence corresponding to the negative control or potential slippery sequence (CT, HIV or $MYC_{673-763}$), and (iii) a second cistron introduced in the (-1) or (+1) reading frame of RLuc that allows to translate the Firefly Luciferase (FLuc) reporter protein when a (-1) or (+1) ribosomal frameshift occurs on the linker sequence. All sequences (CT, HIV, MYC673-763, Renilla and Firefly Luciferases) are available in Table S1. Negative control (CT) and positive slippery control (HIV(-1)) sequences for ribosomal frameshift were designed as previously described (*56*). The potential $MYC_{673-763}$ slippery sequence corresponds to 91-long nucleotide region located upstream the nucleotide sequence encoding the PR3 peptide and between two non-canonical translational start sites in-frame of PR3. For pTRIP_CT, pTRIP_HIV(-1) and pTRIP_MYC(-1), FLuc is -1 reading frame relative to RLuc. For pTRIP_MYC(+1), FLuc is in +1 reading frame relative to RLuc.

<u>Dual-luciferase frameshift assay</u>

Cell lines expressing each bicistronic reporter plasmids were obtained by lentiviral transduction. Using the X-tremGENE HP kit (Roche), lentiviruses were produced by co-transfecting 1.3 x $10^5$ 293T cells with three plasmids (ratio 4:2:4): pCMVdeltaR8.91 and phCMVG-VSVG packaging plasmids (kind gift of Dr. AP Morel (*57*)); and one of the four bicistronic plasmids (pTRIP_CT, pTRIP_HIV(-1), pTRIP_MYC(-1) or pTRIP_MYC(+1)). After 48 h, the supernatant was collected, filtered, supplemented with 5 µg/ml polybrene (Sigma-Aldrich), and inoculated for 10 h on the targeted cells, plated the previous day at 10,000 cells per well in a 6-well plate. After 7 days, stable transduced cells were used for a dual-luciferase assay. 10,000 to 30,000 cells were seeded per well in 96-well plates. After 48 h, Renilla Luciferase (RLuc) and Firefly Luciferase

(FLuc) activities were measured using the Dual-Luciferase Reporter Assay System (Promega) on Spark plate reader (Tecan) according to the manufacturer's protocol. Results were obtained from background-subtracted values of non-transduced cells. Frameshift index corresponds to the FLuc/RLuc ratio.

RT-qPCR analysis

Total RNA was extracted using the RNeasy Plus Mini Kit (Qiagen) according to the manufacturer's protocol. RNAs were quantified using NanoDrop (Thermo Fisher Scientific). cDNA was synthesized using the PrimeScript RT reagent kit (Takara Bio) according to the manufacturer's instructions. Then, qPCR was performed using a LightCycler (Roche) in combination with the LightCycler LC480 SYBR Green I Master (Roche). For *c-MYC* splice variant analysis, 2 x $10^{-15}$ µg of linearized double-stranded DNA coding for *c-MYC* splice variant was used as positive control (synthetized by RD Biotech). *c-MYC* splice variant and all primer sequences used are available in Table S2. For *c-MYC* splice variant analysis, *c-MYC* splice variant relative expression was calculated as: $2^\wedge(-CT_{sample})/2^\wedge(-CT_{positive\ ctrl})$. For the dual-luciferase assay, the relative quantification of Firefly (RQ_FLuc) and Renilla (RQ_RLuc) RNA levels to that of the calibrator was calculated as: 2^-(ΔCTsample - ΔCTcalibrator), where CT is the threshold cycle, ΔCT is (CT target gene – CT GAPDH) and the calibrator is the HMEC_hTERT cell line expressing the CT bicistronic construction. Firefly/Renilla RNA ratio was calculated as: RQ_FLuc/ RQ_RLuc.

PR3-specific $CD8^+$ T cell generation

Healthy donor monocyte-derived dendritic cells (MoDC)/ peripheral blood lymphocytes (PBL) priming was performed as previously described (*42*), with additional modifications**.** MoDCs were maturated overnight (18 h) using 20 ng/ml TNFα (PeproTech) and 40 µg/ml Poly(I:C) (Invivogen) and pulsed in corresponding conditions with MART-1 (2.5 µg/ml) or PR3 peptide (10 µg/ml). MoDC/ PBL were co-cultured in AIM-V (Gibco) with 8% human serum AB (sAB, Eurobio) and 1% PS, in 96-well plates. At day 5, and every 2 to 3 days, medium was replaced with IL-7 (Miltenyi) and IL-15 (Peprotech) supplementation at 10 ng/ml each and cells were split if needed. After 12 days, PR3-dextramer positive and negative $CD8^+$ T cells were sorted using BD FACSAria Cell Sorter (BD Bioscience) and expanded on a feeder as previously described(*42*), with additional modifications. At day 5, and every 2 to 3 days, medium was replaced with 8% sAB, IL-7 and IL-15 supplementation at 20 ng/ml each. After 14 days, PR3-specific $CD8^+$ T cell purity was assessed by dextramer staining. For all experiments, PR3-specific $CD8^+$ T cells with a purity > 90% and autologous dextramer-negative $CD8^+$ T cells were cultured in AIM-V with 8% sAB and 1% PS.

Single-cell TCR sequencing

Single-cell capturing and downstream library constructions of PR3-specific $CD8^+$ T cells were performed using Chromium Next GEM Single Cell 5' Reagent Kits v2 Chemistry Dual Index (10x Genomics, PN1000190, PN1000252, PN1000263) according to the manufacturer's instructions. The constructed scVDJ and scGEX libraries were sequenced on Novaseq 6000 system (Illumina). Clonotype abundance and TCRα/β sequences were analyzed using CellRanger (10x Genomics).

Lentivirus production expressing PR3-specific TCR

Clone 1 PR3-TCRα/β sequences were synthesized and cloned into the pSDY lentiviral backbone (RD Biotech). To promote proper chain pairing, the human variable regions of PR3-TCRα/β were fused to a modified murine constant chain (mTCR) with a second disulfide bond. Lentivirus expressing PR3-TCRα/β were produced in 293T cells. Briefly, 293T cells were seeded at 0.8 x $10^6$ cells per 100-$mm^2$ plate and were transfected 3 days later with the following plasmid mix: psPAX2 : pMDG : pSDY_PR3-TCRαβ plasmid (RD Biotech), at a 1:1.5:3.6 ratio, using the Calcium Phosphate transfection kit (Invitrogen) according to the manufacturer's protocol. The next day, cell media was replaced by DMEM, 1% PS,10mM HEPES (Gibco). Viral supernatants were collected 48 h and 72 h after transfection, pooled and centrifuged for 16h at 3,000g and 4°C. Lentivirus production medium was also centrifuged in the same condition as a control for transduction. Lentivirus or medium were aliquoted and stored at -80°C.

TCR transduction for TCR-T cells generation

PBMCs were incubated with Dynabeads™ Human T-Expander CD3/CD28 (ratio 1:1 beads:$CD3^+$ T cells, Gibco) for 20min before magnetic sorting. Activated $CD3^+$ T cells were cultured for 48 h in X-Vivo15 (Lonza) with 8% sAB,1% PS, IL-7 and IL-15 (10 ng/ml each). Endogenous TCRα/β was deleted using CRISPR-Cas9 technology. Briefly, ribonucleoprotein (RNP) complexes were formed by hybridization of CRISPR RNAs (crRNAs) targeting the TRAC and TRBC (Integrated DNA Technology) designed as previously described (*58*), trans-activating crispr RNA (tracrRNA, 1072533, Integrated DNA Technology), and TrueCut Cas9 protein (Thermo Fisher Scientific). RNP were electroporated into activated $CD3^+$ T cells using Neon™ transfection system (Invitrogen). CRISPR TCRα/β $CD3^+$ T cells were seeded at 1 x $10^5$ cells per well in a previously RetroNectin-coated (1 µg/ml, Takara) 24-well plate and transduced with PR3-TCRα/β lentivirus or lentivirus production media as control (non-transduced condition, NT). After 4 days, cells were collected and the transduction efficacy was evaluated by multiparametric flow analysis. For *in vivo* experiments, $CD8^+$ T cells were sorted using $CD4^+$ depletion (CD4 microbeads, Miltenyi) according to the manufacturer's instructions before incubation with CD3/CD28 beads. For all experiments, TCR-T cells were cultured in X-Vivo15 with 8% sAB and 1% PS.

IFNγ ELISpot

T2 cells were pulsed with a limiting PR3 peptide dilution from $10^{-9}$ to $10^{-13}$ M or irrelevant peptide at $10^{-9}$ M for 5 h at 37°C. After extensive washes, pulsed T2 cells were resuspended in corresponding T cell medium defined above and co-cultured with T cells at 1:10 E:T ratio, on an antibody-coated 96-well PVDF-bottomed plate (Millipore). After 18 h, IFNγ production was detected using Human IFNγ ELISpot Set (Diaclone), following the manufacturer's instructions. Revealed plates were imaged on an ImmunoSpot S6 Ultimate image analyzer and analyzed with the ImmunoSpot analysis software. The peptide concentration required to achieve a half-maximal cytokine response ($EC_{50}$) was determined (GraphPad Prism v10, $R > 0.99$).

T cell immunoassay and cytokine production

Specific cytotoxic activity and activation of T cells were assessed by co-culturing T2 target cells, previously labelled for 13 min at 37°C with CFSE (1/2000, C34554, Thermo Fisher Scientific) and pulsed for 2 h with the PR3 or an irrelevant peptide (10 µg/ml), with PR3-specific or unspecific T cells at E:T ratios indicated in the figures. After 24 h, supernatants were collected to evaluate cytokine production by ELISA (IFNγ and TNFα ELISA, Thermo Fisher Scientific; Granzyme B and IL-2 ELISA, R&D systems) according to the manufacturer's instructions. In parallel, the percentage of CFSE$^{+}$ target cell death and CD137 (4-1BB) expressing T cells were assessed by multiparametric flow cytometry, using viability (Zombie NIR Fixable viability kit, Cat#423106, Biolegend) and cell surface markers such as anti-human CD3 (BV421, 1/40, Cat#306718, Biolegend), anti-human CD8 (APC, 1/100, Cat#344722, Biolegend) and anti-CD137 (PE-Dazzle594, 1/30, Cat#309826, Biolegend). The percentage of lysis corresponds to the cell death normalized against target cells alone, and the percentage of specific lysis to the cell death normalized against target cells pulsed with irrelevant peptide.

T cell cytotoxicity assays

T cell cytotoxic activity against tumor cell lines or normal primary cells was measured in real time using IncuCyte S3 (Sartorius) or xCELLigence RTCA eSight (Agilent). Target cells were seeded onto 96-well plates for IncuCyte or E-Plate VIEW 96 (Agilent) for xCELLigence and cultured at 37°C overnight. According to conditions, a blocking anti–MHC-I antibody (50µg per well; clone W6/32, BioXCell), the isotype control or the PR3 peptide (10 µg/ml per well) were added for 2 h at 37°C. PR3-pulsed cells were cautiously washed with warm medium, and T cells were added in a 2:1 E:T ratio, in the presence of IncuCyte Cytotox Green Dye (1/400, Sartorius).

For IncuCyte experiments, live imaging was performed over 30 h at 37°C. Cell death was calculated as the total number of green stained target cells per well, normalized against the number of target cells per well for mCherry-transduced tumor cells. Maximum killing was established using PR3-pulsed target cells. Lysis was calculated according to the following formula: Lysis (%) = ((T cell-induced target cell death − spontaneous target cell death)/(maximum killing − spontaneous target cell death)) x 100.

For xCELLigence experiments, impedance measurements (cell index) were recorded every 15 min for up to 48 h. The normalized cell index (NCI) was calculated by normalizing the cell index at the time of T cell addition. Percent lysis was calculated using RTCA Software Pro Immunotherapy Module.

TCRαβ-PR3-MHC prediction model

3D models were predicted with AlphaFold2 (*59, 60*). The prediction was run online with ColabFold v1.5.3 (*61*) and the following options: no templates, model=alphafold2_multimer_v3, number of recycling=3, pair mode=unpaired_paired, msa_mode=mmseqs2_uniref_env. Five models were generated and ranked by the multimer composite score (0.8*ipTM +0.2pTM). Only the best model was minimized and kept for further analysis. In addition to the atom coordinates, AlphaFold predicts confidence scores for each model. The first type of confidence score is the predicted value of the local Distance Difference Test (plDDT), a local confidence measure that predicts the quality of the local environment of each residue (fig. S5C). The second type of confidence score is the Predicted Aligned Error (PAE) matrix, that predicts the error in the relative orientation between two regions of a protein or a complex (fig. S5D). Global scores are derived from the PAE matrix: the pTM score is a global quality score and the ipTM is a global quality score restricted to the relative orientation between distinct chains in a complex. The 3D models were analyzed using USCF ChimeraX (*62*).

Amino-acid scanning

HLA-A2$^{+}$ T2 cells were pulsed overnight at 37°C with either the PR3 peptide, irrelevant peptide or variants of PR3 that contain an alanine or glycine substitution at each individual peptide position (10 ng/ml). After extensive washes, pulsed T2 cells were co-cultured with PR3-specific TCR-T cells at an E:T ratio of 10:1. After 24h, IFNγ production was measured using enzyme-linked immunosorbent assay (IFNγ ELISA, Thermo Fisher Scientific). The IFNγ normalized secretion was calculated according to the following formula: $IFN\gamma\ (\%\ max) = ([IFN\gamma_{peptide\text{-}pulsed\ T2}]-[IFN\gamma_{irr\text{-}pulsed\ T2}])/([IFN\gamma_{PR3\text{-}pulsed\ T2}]-[IFN\gamma_{irr\text{-}pulsed\ T2}]) \times 100$. A residue of the PR3 sequence was considered involved in the interaction with the TCR if its substitution resulted in ≥50% loss of function compared to the native peptide sequence for at least 1 replicate, as previously described (*63*). PR3-TCR cross-reactivity screening was performed using ScanProsite to search all UniProtKB/Swiss-Prot database for proteins containing the motif "L-L-x-E-x-T-A-N-L" or "L-x-x-E-x-T-A-N-x". Among ScanProsite hits, HLA-A2 strong binders were assessed using netMHCpan v4.1.

Multiparametric flow cytometry

Dextramer staining was performed on MoDC/PBL priming after a 12-day culture. Cells were washed in PBS + 2% FBS + 2mM EDTA and stained with dextramers (HLA-A*0201/LLLEATANL(PR3)/PE or HLA-A*0201/ELAGIGILTV(MART-1)/PE, Immudex) according to the manufacturer's protocol, in addition to viability (Zombie NIR Fixable viability kit, 1/400, Cat#423106, Biolegend) and cell surface markers such as anti-human CD3 (BV421, 1/40, Cat#300434, Biolegend) and anti-human CD8 (FITC, 1/100, Cat#344704, Biolegend). For tetramer staining, tetramers were produced by conjugate biotinylated HLA-A*02:01 PR3 monomers (P2R Facility, Nantes) with PE- Streptavidin (Cat#405204, Biolegend) or APC-Streptavidin (Cat#405207, Biolegend) for 1 h at room temperature. TCR-T cells were stained with 10 µg/ml tetramer, Zombie NIR Fixable viability, and cell surface markers such as anti-human CD4 (BV605, 1/50, Cat#300556, Biolegend), anti-human CD8 (FITC, 1/100, Cat#344704, Biolegend), anti-mouse TCRβ Chain (BV421, 1/50, Cat#562839, BD Biosciences) and anti-human TCR α/β (APC, 1/50, Cat#306718, Biolegend) for 1 h at 4°C. TILs were stained with 20 µg/ml tetramer, viability (Fixable Viability Stain 780, 1/1000, Cat#565388, BD Biosiences), and cell surface markers such as anti-human CD3 (BUV395, 1/40, Cat#563546, BD Biosiences), anti-human CD8 (BV421, 1/100, Cat#301036, Biolegend). All data were acquired on an LSRFortessa using BD FACSDiva 8.0 software (BD Biosciences) and analyzed using FlowJo software (v10.9.0, Tree Star).

*in vivo* TCR-T cells anti-tumoral effects in chicken embryos

*In vivo* anti-tumoral activity of CD8$^{+}$ PR3-specific TCR-T cells was performed by Oncofactory (Lyon, France). Embryonated eggs were obtained from a local supplier (EARL Les Bruyeres, France) with the sanitary status of laying hens regularly checked by the supplier according to French laws. Development of embryos and *in ovo* xenografts were performed as previously described (*24*), with further modifications. At HH14, embryo's presumptive somitic areas were co-engrafted with previously mixed CFSE-labelled cancer cells (MDA-MB-231 or OVCAR-3) and CD8$^{+}$ PR3-specific or CD8$^{+}$ NT T cells at a 5:1 E:T ratio. After 48h (HH25), embryos were harvested and *in vivo* TCR-T cells anti-tumoral effects was evaluated by assessing the tumor volume using 3D light-sheet imaging. For this analysis, tissues from n=59 chicken embryos (n=33 for MDA-MB-231 and n=26 for OVCAR3 group) were cleared and imaged using whole-mount SPIM imaging with volumetric analyses performed as previously described (*24*). In addition, immunofluorescence on embryo cryosections was performed. Additional chicken embryos (n=6 for MDA-MB-231 and n=6 for OVCAR-3) were harvested and fixed in 4% paraformaldehyde (PFA). Embryos were embedded in 7.5% gelatin- 15% sucrose in PBS to perform 12 µm transverse cryosections. Slides were incubated 20 min with Formol 10%, washed, dried 30 min, and then placed in a Bond RX automated immunostainer (Leica biosystems). Immunofluorescence of $CFSE^{+}$ labeled tumor cells was performed using the OPAL 6-Plex detection kit (AKOYA biosciences). Nuclei were stained with DAPI then mounted using Prolong™ Gold Antifade Reagent (Invitrogen). Slices were scanned using PhenoImager HT software (Akoya bioscience) at 40X, unmixed using the spectral library from INFORM 3.0. Whole slide views for native image were generated using Phenochart and quantitative analysis was performed using HALO (Indica Labs). The results show the number of tumor cells in the slide with the most tumor cells quantified, for each embryo.

Statistical analysis

Data are presented as mean ± s.d. Wilcoxon rank tests were used when appropriate and as indicated. P values were considered statistically significant if P<0.05, with *P<0.05 and ***P<0.001. All statistical values measurements were performed using R v4.3.3 and detailed analysis results are available Table S3.

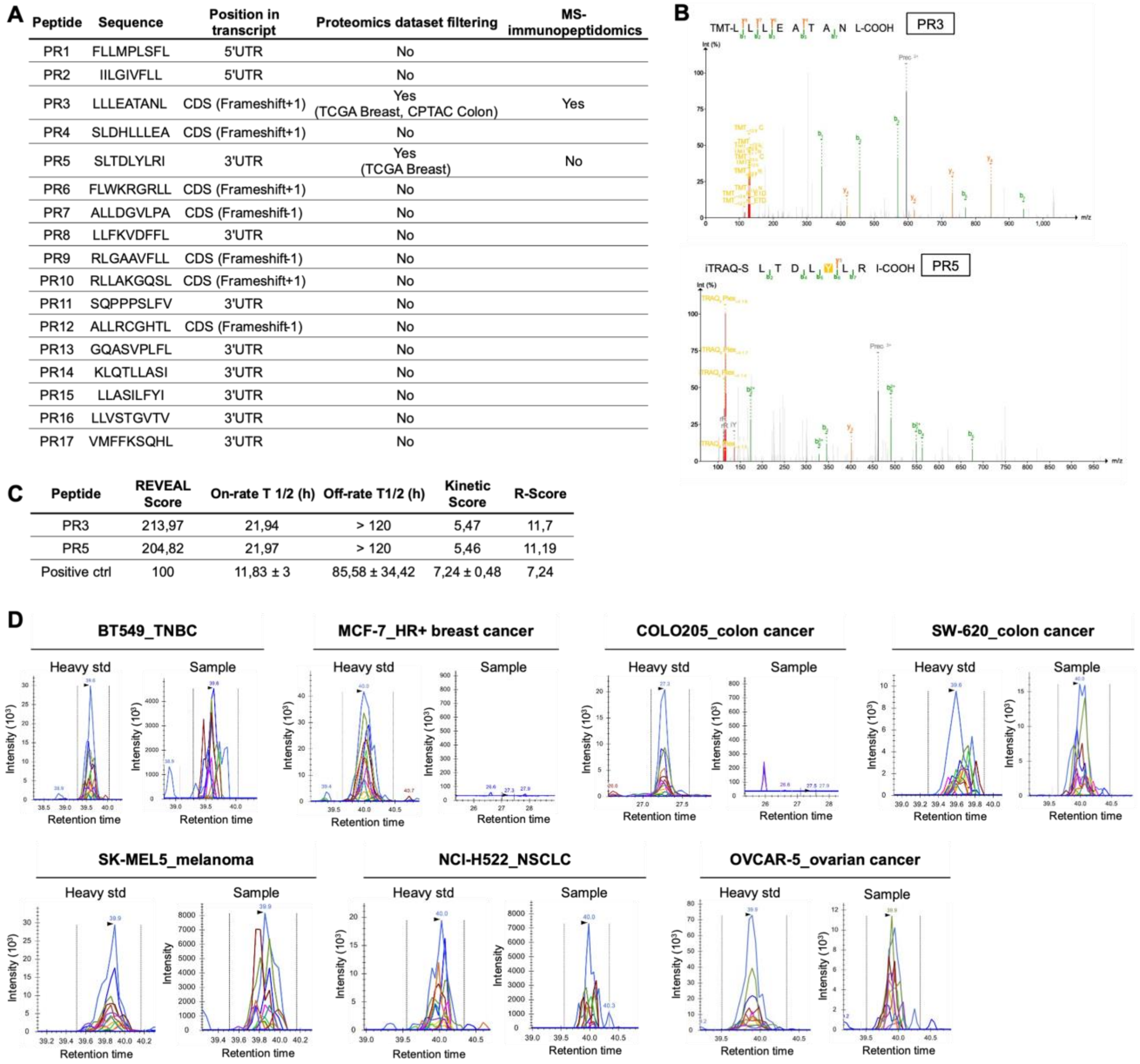


| Peptide | Sequence | Position in transcript | Proteomics dataset filtering | MS-immunopeptidomics |
|---|---|---|---|---|
| PR1 | FLLMPLSFL | 5'UTR | No | |
| PR2 | IILGIVFLL | 5'UTR | No | |
| PR3 | LLLEATANL | CDS (Frameshift+1) | Yes (TCGA Breast, CPTAC Colon) | Yes |
| PR4 | SLDHLLLEA | CDS (Frameshift+1) | No | |
| PR5 | SLTDLYLRI | 3'UTR | Yes (TCGA Breast) | No |
| PR6 | FLWKRGRLL | CDS (Frameshift+1) | No | |
| PR7 | ALLDGVLPA | CDS (Frameshift-1) | No | |
| PR8 | LLFKVDFFL | 3'UTR | No | |
| PR9 | RLGAAVFLL | CDS (Frameshift-1) | No | |
| PR10 | RLLAKGQSL | CDS (Frameshift+1) | No | |
| PR11 | SQPPPSLFV | 3'UTR | No | |
| PR12 | ALLRCGHTL | CDS (Frameshift-1) | No | |
| PR13 | GQASVPLFL | 3'UTR | No | |
| PR14 | KLQTLLASI | 3'UTR | No | |
| PR15 | LLASILFYI | 3'UTR | No | |
| PR16 | LLVSTGVTV | 3'UTR | No | |
| PR17 | VMFFKSQHL | 3'UTR | No | |

| Peptide | REVEAL Score | On-rate T 1/2 (h) | Off-rate T1/2 (h) | Kinetic Score | R-Score |
|---|---|---|---|---|---|
| PR3 | 213,97 | 21,94 | > 120 | 5,47 | 11,7 |
| PR5 | 204,82 | 21,97 | > 120 | 5,46 | 11,19 |
| Positive ctrl | 100 | 11,83 ± 3 | 85,58 ± 34,42 | 7,24 ± 0,48 | 7,24 |

**Fig. S1. Prediction of translation-associated neoepitopes.** (A) Table showing HLA-A2$^{+}$ *c-MYC*-derived neoepitopes predicted using *in silico* model, their position in *c-MYC* transcript, evidence of translation in mass-spectrometry (MS) datasets and HLA-presentation using MS-immunopeptidomics. UTR, untranslated regions; CDS, coding sequence; TCGA, the cancer genome atlas; CPTAC, the clinical proteomic tumor analysis consortium. (B) MS/MS detection of the PR3 (top) and PR5 (bottom) peptides in samples from CPTAC colon cancer (PR3) and TCGA breast cancer (PR5) dataset. MS/MS spectrum is identified by Pepquery analysis ("confident" peptide spectrum match according to Pepquery). Int, intensity; m/z, mass/charge ratio. (C) Table showing binding rate data and kinetic scores for the PR3, PR5 and positive control peptides. REVEAL scores ≥ 45% indicate good binders (see methods). (D) Valid-NEO transitions of the PR3 peptide on chromatograms, using tumor cell lines as samples, and synthetized heavy standard peptides as positive control. TNBC, triple-negative breast cancer; HR, hormone receptor; NSCLC, non-small-cell lung cancer; std, standard.

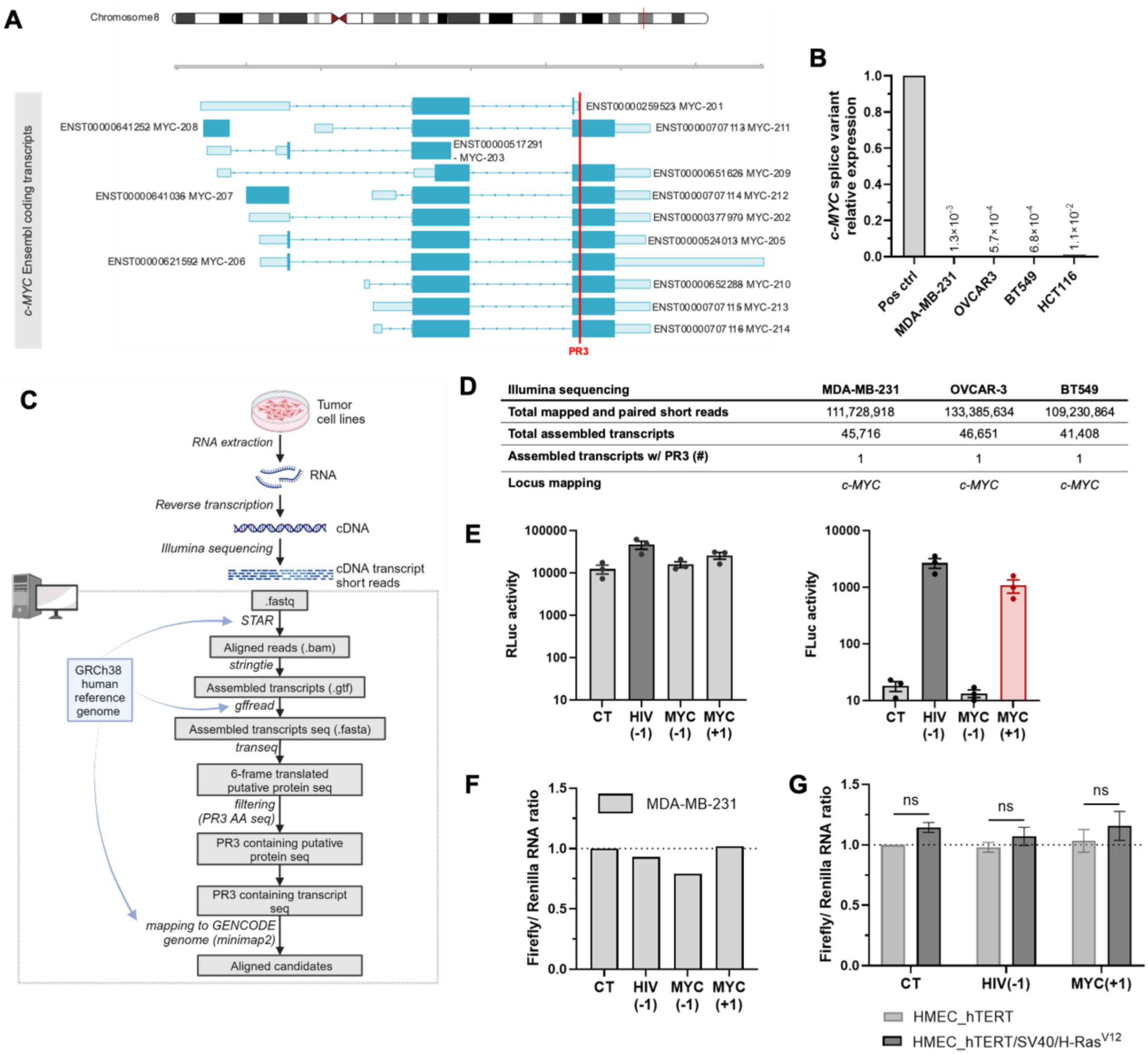


**Fig. S2. Translational origin of PR3.** (A) Overview of aligned *c-MYC* protein-coding transcripts reported in Ensembl. The CDS region of each transcript is represented in dark blue, untranslated regions in light blue, and the position of the PR3 sequence is indicated with a red line. (B) Expression analysis of the putative *c-MYC* splice variant, using RT-qPCR. (C) Schematic overview of the analyses performed using Illumina sequencing. RNA, ribonucleic acid; cDNA, complementary desoxyribonucleic acid; seq., sequence; AA, amino acid. (D) Table showing the locus mapping of the Illumina short read-assembled transcripts containing the PR3 sequence, in MDA-MB-231, OVCAR-3 and BT549 cells. w/, with; #, number. (E) Levels of Renilla (left) and Firefly (right) activities in MDA-MB-231 cells expressing CT, HIV(-1), MYC(-1) or MYC(+1) construct. Results are mean ± s.d. of n=3 independent experiments represented by each dot. (F and G) Ratios of Firefly and Renilla relative quantitation in MDA-MB-231 (F), in HMEC_hTERT and HMEC_hTERT/SV40/H-RAS$^{V12}$ cells (G), expressing CT, HIV(-1), MYC(-1) or MYC(+1) construct, as evaluated by RT-qPCR. Results are mean ± s.d. of n=3 (G) independent experiments. CT, control; HIV, human immunodeficiency viruses; ns, not significant, Wilcoxon test.

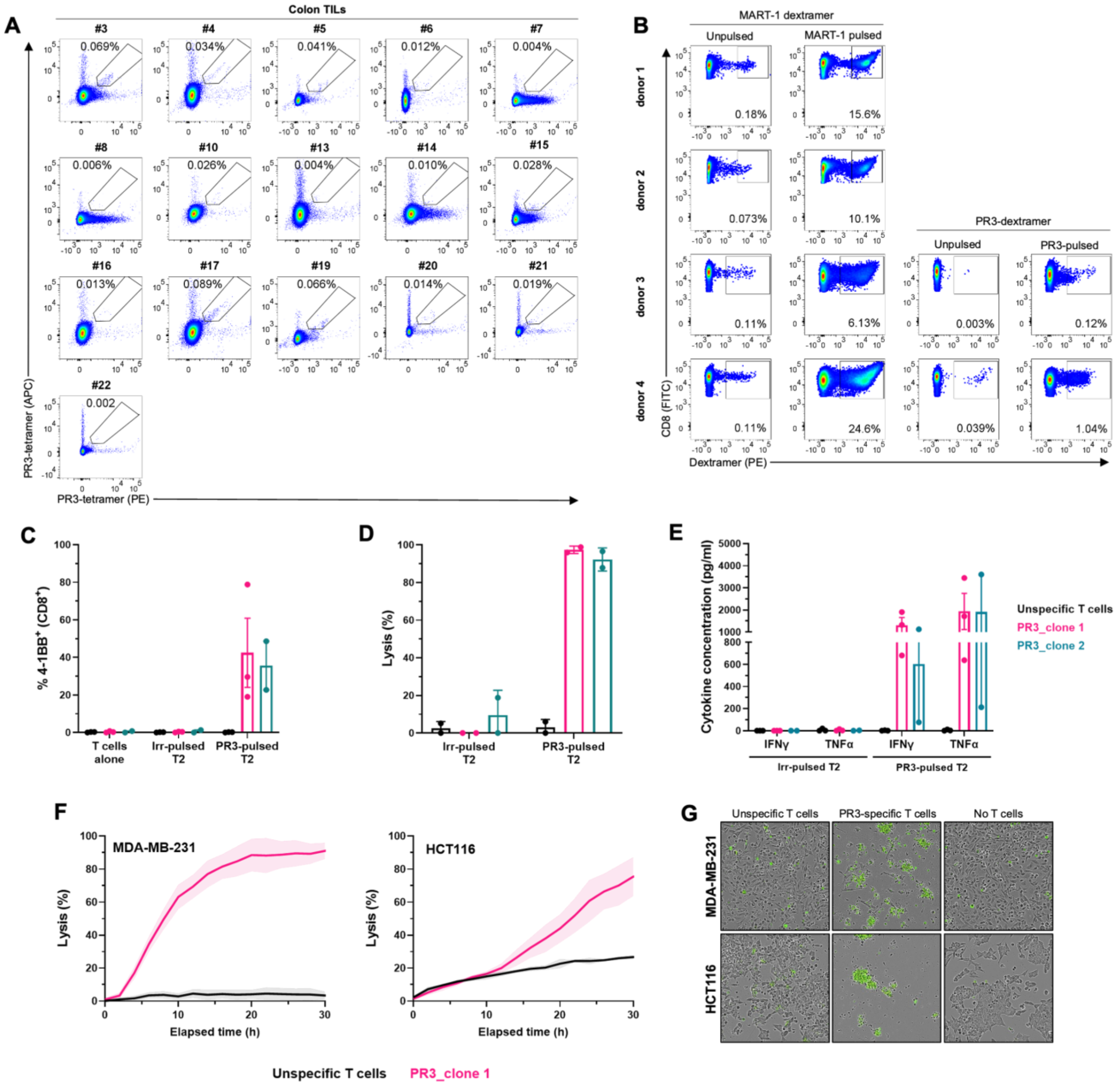


**Fig. S3. Functionality of PR3-specific CD8$^+$ T cells.** (A) Flow cytometry plots and quantification of double PE-/APC-PR3 tetramer staining of TILs from colon cancer samples gated on CD8$^+$ T cells. #, number. (B) Flow cytometry plots and quantification of dextramer staining (MART-1 positive control or PR3) of CD8$^+$ T cells from 4 HLA-A2$^+$ healthy donors, after 12 days of co-culture with unpulsed or peptide-pulsed (MART-1 or PR3) MoDCs. (C to E) PR3-specific CD8$^+$ T cells from clone 1 (pink), clone 2 (blue) or unspecific CD8$^+$ T cells (black) were co-cultured with T2 cells pulsed with PR3 or an irrelevant peptide for 24h at 10:1 effector to target ratio. The percentage of CD137 (4-1BB) gated on CD8$^+$ T cells (C) and lysis of T2 target (D) were determined by flow cytometry. IFNγ and TNFα cytokine secretions (E) were determined by ELISA. Results are mean ± s.d. of n=3 (PR3_clone 1 and unspecific T cells) or n=2 (PR3_clone 2) (C and E) and n=2 (D) independent experiments represented by each dot. Irr, irrelevant. (F) Tumor cell death induced by PR3-specific CD8$^+$ T cells from clone 1 (PR3_clone 1) at 2:1 effector to target cell ratio. Results are ± s.d. of technical triplicate. Data shown are representative of n=4 (MDA-MB-231) or n=2 (HCT116) independent experiments. (G) Representative 10X images at 30h of co-culture presented in (H) and tumor cells alone. Dead cells are depicted in green.

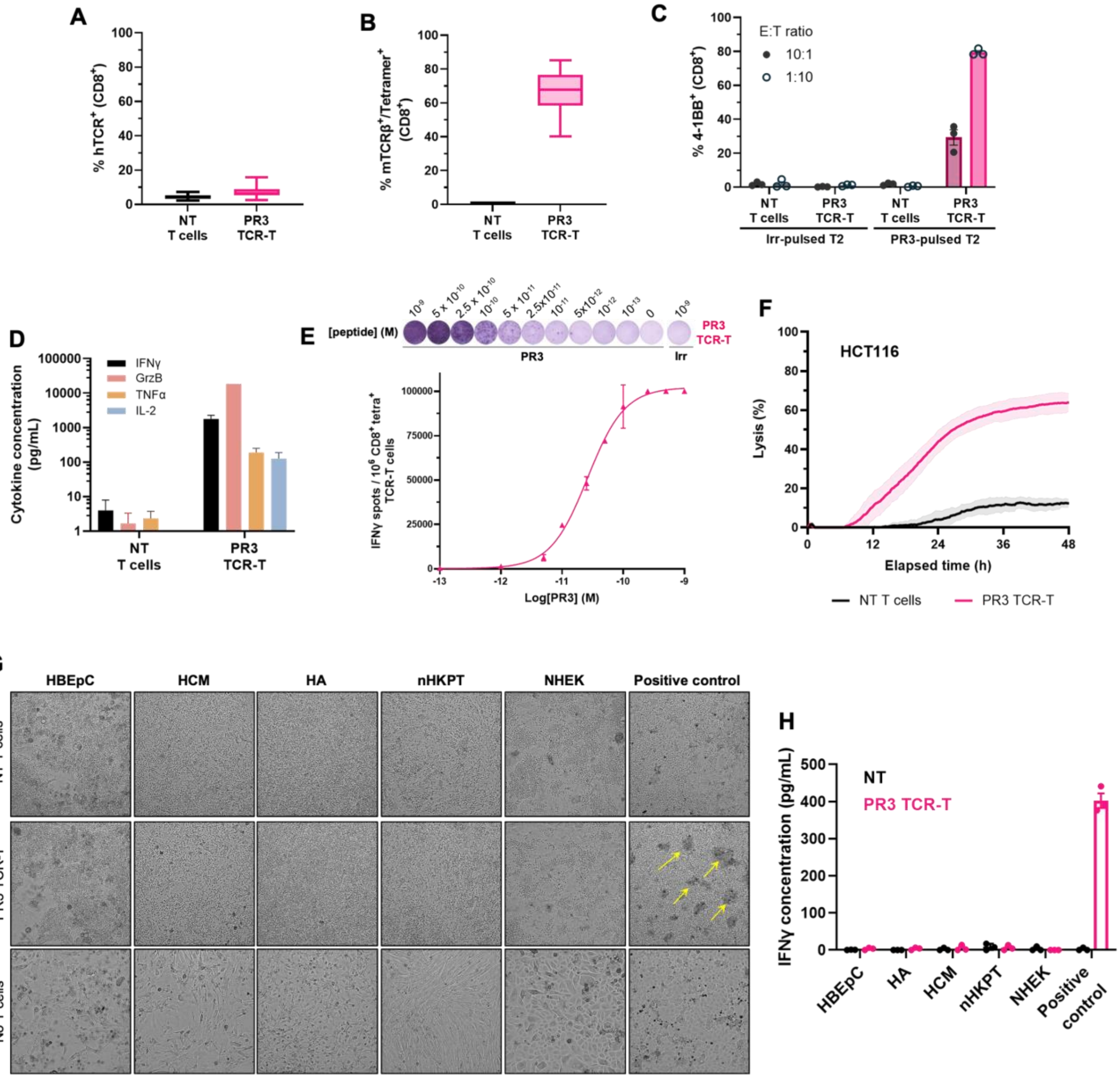


**Fig. S4. Functionality and specificity of PR3-specific TCR-T cells.** (A and B) Box plots showing the percentage of hTCR positive (A) and mTCRβ/PR3-tetramer double positive (B) staining of non-transduced (NT) T cells or PR3-specific TCR-T cells (PR3 TCR-T) gated on $CD8^+$ T cells. Results are mean ± min to max of n=24 (NT) and n=32 (PR3 TCR-T) independent experiments. hTCR, human-TCR. (C) Percentage of CD137 (4-1BB) gated on $CD8^+$ T cells and determined by flow cytometry of NT T cells or PR3 TCR-T co-cultured for 24h with T2 cells pulsed with PR3 or irrelevant peptide, at 10:1 or 1:10 effector to target ratios. E:T: effector to target. (D) Cytokine secretion of NT T cells or PR3 TCR-T after 24h of co-culture with PR3-pulsed T2 cells, normalized against T cells co-cultured with irrelevant pulsed-T2 cells, at 10:1 E:T ratio. GrzB, granzyme-b. Results are mean ± s.d of n=3 (C and D) independent experiments. (E) Representative IFNγ ELISpot images (top), quantification and nonlinear fit curves of IFNγ spot count per $1 \times 10^6$ T cells (bottom) for PR3-specific TCR-T cell (PR3 TCR-T). Results are mean ±

s.d. of technical duplicates. Data shown are representative of n=9 independent experiments. M, molar mass; [ ], concentration; irr, irrelevant. (F) Specific T cell-induced tumor cell death of HCT116 cells at 2:1 E:T ratio. Results are ± s.d. of technical triplicate. (G) Representative 10X images at 48h of co-culture presented in Fig. 4E, with yellow arrows showing T cell clustering associated with target cell death. (H) IFNγ secretion of co-culture presented in Fig. 4E at 48h, determined by ELISA. Results are mean ± s.d. of n=3 independent experiments.

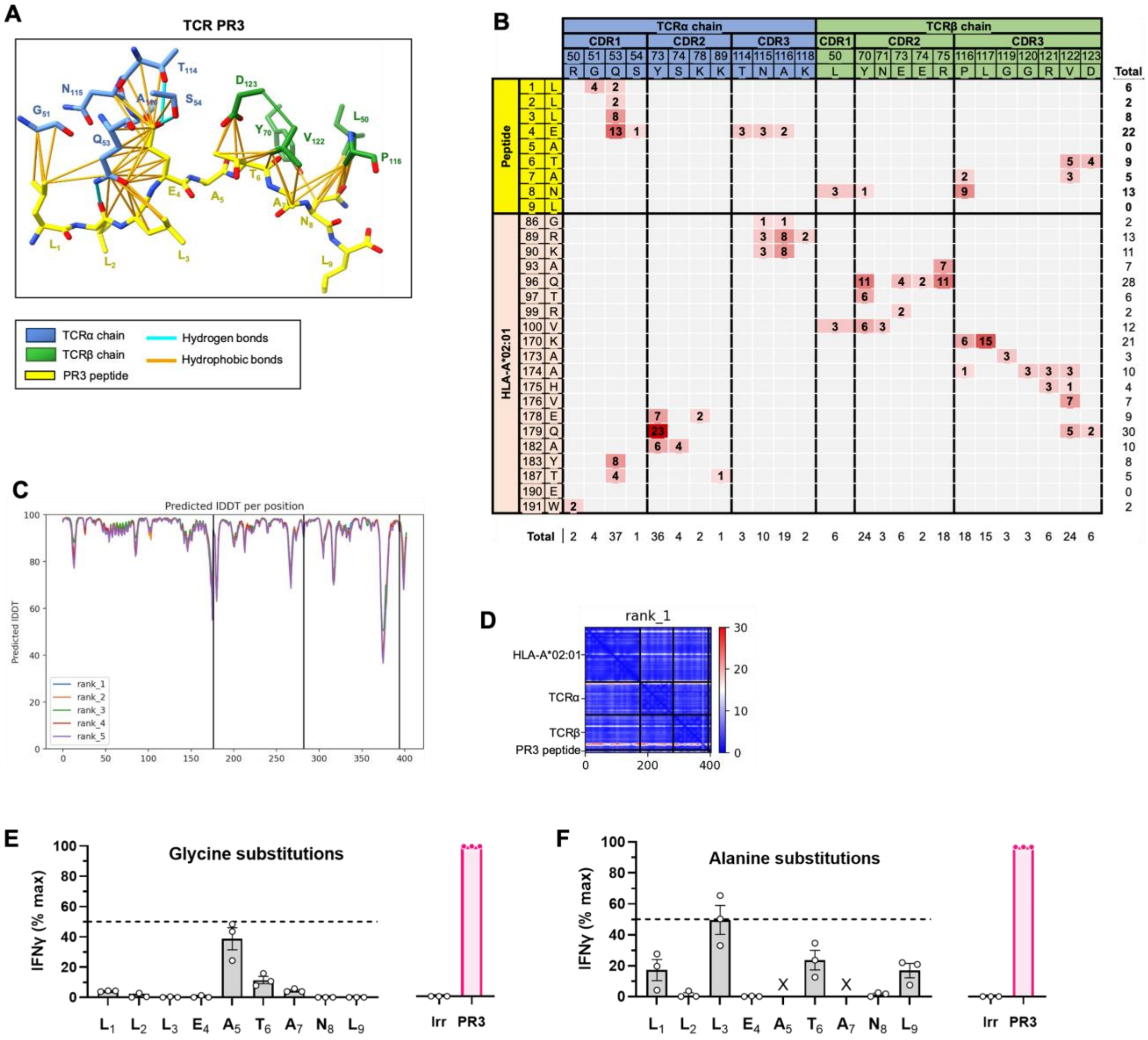


**Fig. S5. TCR modeling and analysis of specificity**. (A) Contacts between the PR3 peptide and the CDR loops of the PR3-specific TCR (TCR PR3) in the AlphaFold2 (AF2) model. The PR3 peptide is depicted in yellow, HLA*02:01 in beige, TCRα in blue and TCRβ in green. The hydrophobic contacts are indicated by orange lines and the hydrogen bounds by blue lines. CDR, complementarity-determining region. (B) Contacts between PR3-HLA*02:01 and TCR in the AF2 model. Atomics contacts are defined using a 5 Å cutoff and the number of contacts is summed on a per-residue basis. Colors correspond to the number of atomic contacts. (C and D) Quality of the AF2 model, with predicted lDDT scores of the 5 models predicted by AF2. (C) and PAE matrix of the best model (ipTM score = 0.9) (D). IDDT, local distance difference test; PAE, predicted aligned error; ipTM, interface predicted template modelling. (E and F) Glycine (E) and alanine (F) scanning. IFNγ secretion of PR3-specific TCR-T cells after 24h of co-culture with T2 cells pulsed with 10 ng/ml glycine (E) or alanine-substituted (F) peptides. X-axis shows each native amino-acid and position of the PR3 peptide sequence, substituted by glycine (E) or alanine (F). Results are mean ± s.d. percent maximum response relative to PR3 peptide and normalized against irrelevant peptide of n=3 independent experiments. ‘x’ indicates positions not amenable to substitution.

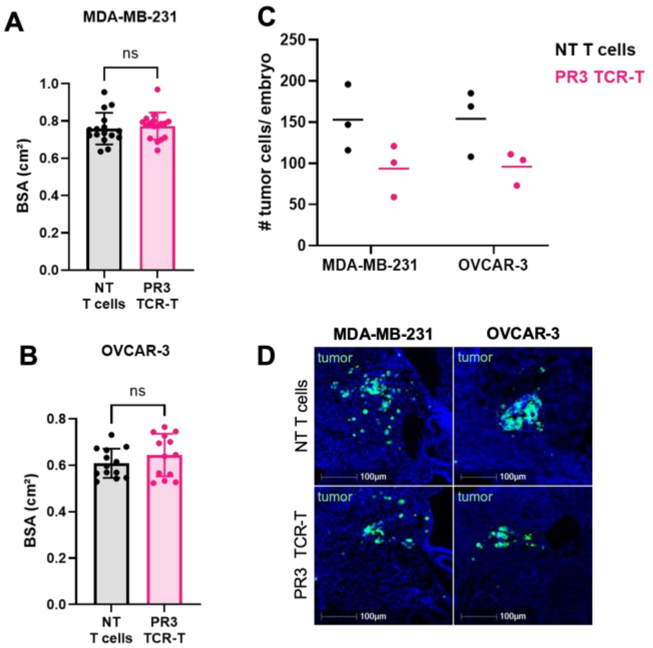


**Fig. S6. *In vivo* experiment quality controls.** (A and B) Quantification of MDA-MB-231 (A) or OVCAR-3 (B) chicken embryo body surface areas (BSA), after 48h co-engraftment with $CD8^+$ non-transduced (NT) T cells or PR3 TCR-T at 5:1 effector to target ratio. Results are mean ± s.d. of 33 embryos (NT T cells=16; PR3 TCR-T=17) (A) and 26 embryos (NT T cells=13; PR3 TCR-T=13) (B) represented by each dot. ns, not significant, Wilcoxon tests. NT, non-transduced**.** (C and D) Immunofluorescence quantification of the number of MDA-MB-231 and OVCAR-3 tumor cells per embryo (n=12, 3 embryos per group) (C) and representative images (D), after 48h co-engraftment with $CD8^+$ NT T cells or PR3 TCR-T at 5:1 effector to target cell ratio. #, number.

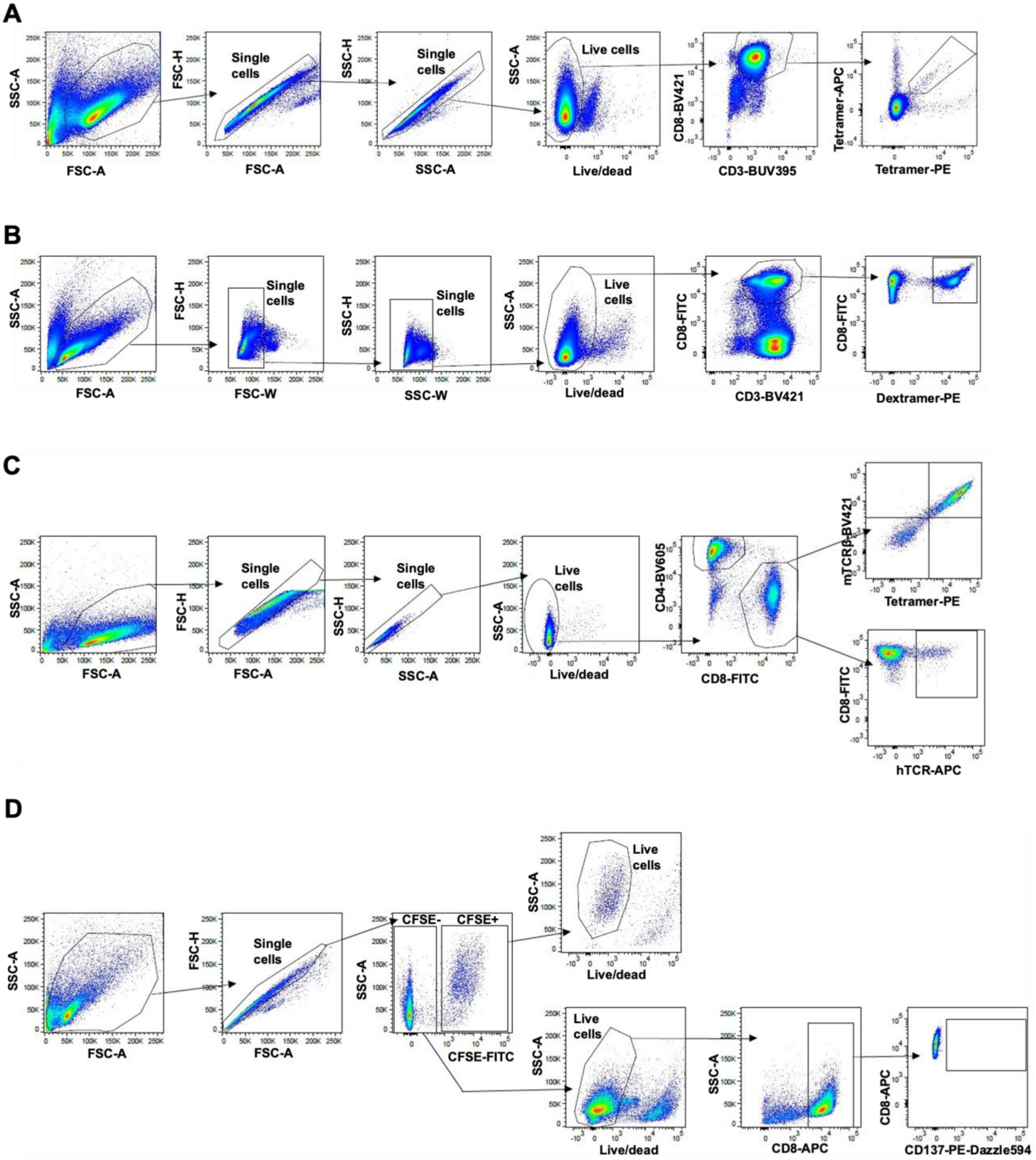


**Fig. S7. Gating strategies for flow cytometry analysis.** (A) The gating strategy of Fig. 3,A and B, and fig. S3A. (B) The gating strategy of Fig. 3, D and E, and fig. S3D. (C) The gating strategy of Fig. 4A and fig. S4, A and B. (D) The gating strategy of Fig. 4D and fig. S3, C and D, and fig. S4C.

| Construct | Sequence (5’ - 3’) |
|---|---|
| CT | ATGTCGACGGATCCTTCAACTTCCCTGAGCTCG |
| HIV(-1) | GATCCTTTTTTAGGGAAGATCTGGCCTTCCCACAAGGGAAGGCCAGGGAATTTTCTTCA<br>GAGCAGACCAGAGCCAACAGCCGCACCGAGCT |
| $MYC_{673\text{-}763}$ | CAAGACTCCAGCGCCTTCTCTCCGTCCTCGGATTCTCTGCTCTCCTCGACGGAGTCCTCC<br>CCGCAGGGCAGCCCCGAGCCCCTGGTGCTCC |
| Renilla Luciferase | ATGACTTCGAAAGTTTATGATCCAGAACAAAGGAAACGGATGATAACTGGTCCGCAGT<br>GGTGGGCCAGATGTAAACAAATGAATGTTCTTGATTCATTTATTAATTATTATGATTCAG<br>AAAAACATGCAGAAAATGCTGTTATTTTTTTACATGGTAACGCGGCCTCTTCTTATTTAT<br>GGCGACATGTTGTGCCACATATTGAGCCAGTAGCGCGGTGTATTATACCAGACCTTATT<br>GGTATGGGCAAATCAGGCAAATCTGGTAATGGTTCTTATAGGTTACTTGATCATTACAA<br>ATATCTTACTGCATGGTTTGAACTTCTTAATTTACCAAAGAAGATCATTTTTGTCGGCCA<br>TGATTGGGGTGCTTGTTTGGCATTTCATTATAGCTATGAGCATCAAGATAAGATCAAAG<br>CAATAGTTCACGCTGAAAGTGTAGTAGATGTGATTGAATCATGGGATGAATGGCCTGAT<br>ATTGAAGAAGATATTGCGTTGATCAAATCTGAAGAAGGAGAAAAAATGGTTTTGGAGA<br>ATAACTTCTTCGTGGAAACCATGTTGCCATCAAAAATCATGAGAAAGTTAGAACCAGAA<br>GAATTTGCAGCATATCTTGAACCATTCAAAGAGAAAGGTGAAGTTCGTCGTCCAACATT<br>ATCATGGCCTCGTGAAATCCCGTTAGTAAAAGGTGGTAAACCTGACGTTGTACAAATTG<br>TTAGGAATTATAATGCTTATCTACGTGCAAGTGATGATTTACCAAAAATGTTTATTGAAT<br>CGGACCCAGGATTCTTTTCCAATGCTATTGTTGAAGGTGCCAAGAAGTTTCCTAATACT<br>GAATTTGTCAAAGTAAAAGGTCTTCATTTTTCGCAAGAAGATGCACCTGATGAAATGGG<br>AAAATATATCAAATCGTTCGTTGAGCGAGTTCTCAAAAATGAACAA |
| Firefly Luciferase | GAAGACGCCAAAAACATAAAGAAAGGCCCGGCGCCATTCTATCCGCTGGAAGATGGAA<br>CCGCTGGAGAGCAACTGCATAAGGCTATGAAGAGATACGCCCTGGTTCCTGGAACAAT<br>TGCTTTTACAGATGCACATATCGAGGTGGACATCACTTACGCTGAGTACTTCGAAATGT<br>CCGTTCGGTTGGCAGAAGCTATGAAACGATATGGGCTGAATACAAATCACAGAATCGT<br>CGTATGCAGTGAAAACTCTCTTCAATTCTTTATGCCGGTGTTGGGCGCGTTATTTATCGG<br>AGTTGCAGTTGCGCCCGCGAACGACATTTATAATGAACGTGAATTGCTCAACAGTATGG<br>GCATTTCGCAGCCTACCGTGGTGTTCGTTTCCAAAAAGGGGTTGCAAAAAATTTTGAAC<br>GTGCAAAAAAAGCTCCCAATCATCCAAAAAATTATTATCATGGATTCTAAAACGGATTA<br>CCAGGGATTTCAGTCGATGTACACGTTCGTCACATCTCATCTACCTCCCGGTTTTAATGA<br>ATACGATTTTGTGCCAGAGTCCTTCGATAGGGACAAGACAATTGCACTGATCATGAACT<br>CCTCTGGATCTACTGGTCTGCCTAAAGGTGTCGCTCTGCCTCATAGAACTGCCTGCGTGA<br>GATTCTCGCATGCCAGAGATCCTATTTTTGGCAATCAAATCATTCCGGATACTGCGATTT<br>TAAGTGTTGTTCCATTCCATCACGGTTTTGGAATGTTTACTACACTCGGATATTTGATAT<br>GTGGATTTCGAGTCGTCTTAATGTATAGATTTGAAGAAGAGCTGTTTCTGAGGAGCCTT<br>CAGGATTACAAGATTCAAAGTGCGCTGCTGGTGCCAACCCTATTCTCCTTCTTCGCCAA<br>AAGCACTCTGATTGACAAATACGATTTATCTAATTTACACGAAATTGCTTCTGGTGGCG<br>CTCCCCTCTCTAAGGAAGTCGGGGAAGCGGTTGCCAAGAGGTTCCATCTGCCAGGTATC<br>AGGCAAGGATATGGGCTCACTGAGACTACATCAGCTATTCTGATTACACCCGAGGGGG<br>ATGATAAACCGGGCGCGGTCGGTAAAGTTGTTCCATTTTTTGAAGCGAAGGTTGTGGAT<br>CTGGATACCGGGAAAACGCTGGGCGTTAATCAAAGAGGCGAACTGTGTGTGAGAGGTC<br>CTATGATTATGTCCGGTTATGTAAACAATCCGGAAGCGACCAACGCCTTGATTGACAAG<br>GATGGATGGCTACATTCTGGAGACATAGCTTACTGGGACGAAGACGAACACTTCTTCAT<br>CGTTGACCGCCTGAAGTCTCTGATTAAGTACAAAGGCTATCAGGTGGCTCCCGCTGAAT<br>TGGAATCCATCTTGCTCCAACACCCCAACATCTTCGACGCAGGTGTCGCAGGTCTTCCC<br>GACGATGACGCCGGTGAACTTCCCGCCGCCGTTGTTGTTTTGGAGCACGGAAAGACGAT<br>GACGGAAAAAGAGATCGTGGATTACGTCGCCAGTCAAGTAACAACCGCGAAAAAGTTG<br>CGCGGAGGAGTTGTGTTTGTGGACGAAGTACCGAAAGGTCTTACCGGAAAACTCGACG<br>CAAGAAAAATCAGAGAGATCCTCATAAAGGCCAAGAAGGGCGGAAAGATCGCCGTGT<br>AA |

**Table S1. Sequences of bicistronic reporter constructs.**

| Primer sequences | | |
|---|---|---|
| **Gene name** | **Forward sequence** | **Reverse sequence** |
| **Renilla** | AACGCGGCCTCTTCTTATTT | ACCAGATTTGCCTGATTTGC |
| **Firefly** | AACACCCCAACATCTTCGAC | TTTTCCGTCATCGTCTTTCC |
| ***c-MYC* splice variant** | CGGGTAGTGGAAAACCAGAGG | CTCCAGCAGAAGGTGATCCAG |
| **GAPDH** | AGCCACATCGCTCAGACAC | GCCCAATACGACCAAATCC |

| DNA control for *c-MYC* splice variant quantification by RT-qPCR | |
|---|---|
| **Gene** | **Sequence (5' - 3')** |
| ***c-MYC* splice variant** | CTGGATTTTTTTCGGGTAGTGGAAAACCAGAGGAGGAACAAGAAGA<br>TGAGGAAGAAATCGATGTTGTTTCTGTGGAAAAGAGGCAGGCTCCT<br>GGCAAAAGGTCAGAGTCTGGATCACCTTCTGCTGGAGGCCACAGCA<br>AACCTCCTCACAGCCCACTGGTCCTCAAGAGGTGCCACGTCTCCAC<br>ACATCAGCACAACTACGCAGCGCCTCCCTCCACTCGGAAGGACTAT<br>CCTGCTGCCAAGAGGGTCAAGTTGGACAGTGTCAGAGTCCTGA |

**Table S2. Primers and control sequence used in the RT-qPCR.**

| Statistical analysis of Fig. 2E | |
|---|---|
| CT | Wilcoxon rank sum exact test<br>data: Frameshift index<br>W = 1, p-value = 0.05714<br>alternative hypothesis: true location shift is not equal to 0 |
| HIV(-1) | Wilcoxon rank sum exact test<br>data: Frameshift index<br>W = 0, p-value = 0.02857<br>alternative hypothesis: true location shift is not equal to 0 |
| MYC(+1) | Wilcoxon rank sum exact test<br>data: Frameshift index<br>W = 0, p-value = 0.02857<br>alternative hypothesis: true location shift is not equal to 0 |

| Statistical analysis of fig. S2G | |
|---|---|
| CT | Wilcoxon rank sum test with continuity correction<br>data: Firefly/ Renilla RNA ratio<br>W = 0, p-value = 0.0636<br>alternative hypothesis: true location shift is not equal to 0 |
| HIV(-1) | Wilcoxon rank sum test with continuity correction<br>data: Firefly/ Renilla RNA ratio<br>W = 2, p-value = 0.3758<br>alternative hypothesis: true location shift is not equal to 0 |
| MYC(+1) | Wilcoxon rank sum exact test<br>data: Firefly/ Renilla RNA ratio<br>W = 3, p-value = 0.7<br>alternative hypothesis: true location shift is not equal to 0 |

| Statistical analysis of Fig. 5B | |
|---|---|
| MDA-MB-231 | Wilcoxon rank sum test with continuity correction<br>data: Normalized tumor volume<br>W = 206, p-value = 0.01229<br>alternative hypothesis: true location shift is not equal to 0 |

| Statistical analysis of Fig. 5C | |
|---|---|
| OVCAR-3 | Wilcoxon rank sum exact test<br>data: Normalized tumor volume<br>W = 151, p-value = 0.0002971<br>alternative hypothesis: true location shift is not equal to 0 |

| Statistical analysis of fig. S5A | |
|---|---|
| MDA-MB-231 | Wilcoxon rank sum test with continuity correction<br>data: BSA<br>W = 109.5, p-value = 0.3489<br>alternative hypothesis: true location shift is not equal to 0 |

| Statistical analysis of fig. S5B | |
|---|---|
| OVCAR-3 | Wilcoxon rank sum exact test<br>data: BSA<br>W = 68, p-value = 0.4184<br>alternative hypothesis: true location shift is not equal to 0 |

**Table S3. Statistical analyses.**
